# A TROP2/Claudin Program Mediates Immune Exclusion to Impede Checkpoint Blockade in Breast Cancer

**DOI:** 10.1101/2024.12.02.626446

**Authors:** Bogang Wu, Win Thant, Elena Bitman, Ting Liu, Jie Liu, Eleftherios I. Paschalis, Bidish K. Patel, Cole Nawrocki, Katherine H. Xu, Linda T. Nieman, David T. Ting, Bruna de Gois Macedo, Yang Cheng, Kevin Jiang, Fengfei Sun, Nayana Thimmiah, Sheng Sun, Rachel O. Abelman, Veerle I. Bossuyt, Steven J. Isakoff, Laura M. Spring, Aditya Bardia, Leif W. Ellisen

**Affiliations:** Massachusetts General Hospital Cancer Center, Harvard Medical School, Boston, MA, 02114, USA; Department of Immunology, Mayo Clinic, Phoenix, AZ, 85054, USA; Massachusetts Eye and Ear, Harvard Medical School, Boston, MA, 02114, USA; Division of Gastrointestinal and Oncologic Surgery, Department of Surgery, Massachusetts General Hospital, Harvard Medical School, MA, 02114, USA; Ludwig Center at Harvard, Boston, MA 02115, USA

**Keywords:** TROP2, Antibody-Drug Conjugate, sacituzumab govitecan, tight junction, triple negative breast cancer, PD-1, antitumor immunity, tumor immune microenvironment, immune checkpoint blockade, immune exclusion

## Abstract

**Background:** Immune exclusion inhibits anti-tumor immunity and response to immunotherapy, but its mechanisms remain poorly defined. In triple-negative breast cancer (TNBC), an aggressive and generally immune-rich subtype, an immune-cold microenvironment predicts poor prognosis due to a limited response to chemotherapy and immune checkpoint inhibitors. This study aimed to identify mechanisms regulating immune infiltration in TNBC.

**Methods:** We performed spatial transcriptomic analysis comparing immune-enriched versus immune-cold treatment-naïve TNBCs. Functional analyses, including loss-of-function and reconstitution experiments, were conducted to investigate the role of Trophoblast Cell-Surface Antigen 2 (TROP2), a key target of anti-cancer Antibody Drug Conjugates (ADCs), in promoting TNBC progression. A humanized TROP2 syngeneic TNBC model was used to assess the effects of TROP2-targeting in combination with anti-PD1 therapy. Additionally, data from patients treated with immune checkpoint blockade were used to test hypotheses from the preclinical findings.

**Results:** We reveal that TROP2 controls barrier-mediated immune exclusion in TNBC through Claudin 7 association and tight junction regulation. TROP2 expression is inversely correlated with T cell infiltration and predicts poor outcomes in TNBC. We demonstrate that TROP2 is sufficient to drive tumor progression in vivo in a CD8 T cell-dependent manner, while its loss deregulates expression and localization of multiple tight junction proteins, enabling T cell infiltration. We show that TROP2 targeting via hRS7, the antibody component of the ADC Sacituzumab govitecan (SG), enhances the anti-PD1 response and improves T cell accessibility and effector function. Correspondingly, TROP2 expression is highly associated with lack of response to anti-PD1 therapy in human breast cancer.

**Conclusions:** This study defines a new mechanism of barrier-mediated immune exclusion in cancer controlled by TROP2-dependent tight junctions. This mechanism drives tumor progression but can be targeted via TROP2-directed therapy to activate anti-tumor immunity and enhance immunotherapy response.

**What is already known on this topic:** Barrier-mediated immune exclusion is emerging as a mechanism of immune evasion in cancer such as triple-negative breast cancer (TNBC), but its molecular underpinnings remain poorly understood. TROP2 is a surface glycoprotein targeted by antibody-drug conjugates such as Sacituzumab govitecan, yet its functional role beyond drug delivery has been unclear.

**What this study adds:** This study identifies TROP2 as a key regulator of tight junction-mediated immune exclusion in TNBC, independent of its intracellular signaling function. TROP2 promotes an immune-cold tumor microenvironment by enforcing mechanical barriers that limit T cell infiltration.

**How this study might affect research, practice or policy:** These findings establish TROP2 as a functional driver of immune evasion and provide a mechanistic rationale for combining TROP2-targeting therapies with immune checkpoint inhibitors to overcome resistance in immune-excluded tumors.

## INTRODUCTION

Mechanical barriers are a major self-defense mechanism within normal tissues. They exclude invading pathogens, maintain tissue integrity, and precisely regulate permeability to excessive immune infiltration that can damage healthy tissues (1, 2). These barrier functions are co-opted in diverse cancers, where emerging data suggest that extracellular matrix and intercellular junctions exclude immune cells and bioactive molecules as a mechanism of immune evasion (3–5). Such barrier mechanisms are also recognized as an impediment to immunotherapy response (6–9). While barrier-mediated immune exclusion has been studied extensively in normal immune-privileged organs (10–12), a detailed understanding of its role in anti-tumor immunity is lacking.

Triple negative breast cancer (TNBC), defined by the absence of estrogen receptor (ER) and progesterone receptor (PR) expression and HER2 amplification, is a highly heterogeneous and poor-prognosis breast cancer subset. Extensive lymphocyte infiltration is common in TNBC, and tumor-infiltrating lymphocytes are an important clinical predictor of response to chemotherapy. Immune-rich TNBCs, particularly those demonstrating PD-L1 staining on tumor and immune cells, also respond to immune checkpoint blockade (ICB). Conversely, immune-cold TNBCs are generally resistant to both chemotherapy and ICB. Thus, these tumors represent a particular therapeutic challenge as they generally lack effective treatment options.

To reveal mechanisms controlling immune infiltration in TNBC, we carried out an unbiased spatial transcriptomic analysis of treatment-naïve TNBCs, comparing gene expression programs in tumor cells from immune-enriched versus immune-cold tumors. Complementary analyses of multiple TNBC datasets converged on Trophoblast cell surface antigen 2 (TROP2) expression as a shared feature of immune-cold and ICB-resistant tumors. TROP2 is a transmembrane glycoprotein expressed on the cell surface in select epithelial tissues. TROP2 is also overexpressed in a variety of carcinomas including TNBC (13, 14). The potential contribution of TROP2 in cancer remains controversial, however. TROP2 overexpression has been implicated in a variety of tumor phenotypes including proliferation, survival, invasion and stem cell character, while other work suggests a contribution of TROP2 loss to tumorigenic cell signaling (15, 16). Encoded by *TACSTD2*, TROP2 is a paralog of EpCAM and is comprised of extracellular, transmembrane and intracellular domains. Diverse signaling pathways that promote cancer phenotypes are reported to be activated by TROP2, including PI3K/AKT (17), MAPK/ERK (18), JAK/STAT (19), and β-catenin (20), in multiple cases attributed to signaling via the 26-amino acid intracellular domain. In contrast, loss of TROP2 function is observed in human gelatinous drop-like corneal dystrophy (GDLD), an inherited disease characterized by increased corneal basement membrane permeability (21–26). Thus, substantial uncertainty exists regarding the critical functions and mechanisms of TROP2 in cancer.

TROP2 is now a major focus for cancer therapy, as it is the target of multiple recently-developed antibody-drug conjugates (ADCs), including Sacituzumab govitecan (SG). SG was the first ADC approved by the FDA for treatment of TNBC and combines a TROP2-targeted antibody with a topoisomerase 1 (TOP1) inhibitor as cytotoxic payload (27–29). As a single-agent, SG improves both progression-free and overall survival compared to standard chemotherapy for patients with metastatic TNBC (28, 29). Notably, it remains unclear whether targeting TROP2 via SG and other ADCs has therapeutic relevance other than as a TOP1 inhibitor delivery mechanism. Addressing the contribution of TROP2 in this context will likely inform future therapeutic combinations incorporating ADCs, including with immune checkpoint inhibitors.

Here, through in vivo loss- and gain-of-function approaches we reveal a specific role for TROP2 in barrier-mediated immune exclusion that drives tumor progression in TNBC. We find that TROP2 is required for tight junction barrier integrity in TNBC, independent of its intracellular signaling domain, thereby enforcing an immune-poor microenvironment and conferring a poor response to immune checkpoint inhibition. This effect can be reversed via TROP2 targeting to enhance response to anti-PD1 therapy in vivo.

## RESULTS

To define molecular underpinnings of immune-hot versus immune-cold TNBC, we performed spatial transcriptomic analysis on 88 regions of interest (ROIs) from eight treatment-naïve human TNBC specimens (Fig. 1A). Slides were stained for cytotoxic T cells (CD8), tumor epithelial cells (pan-CK), fibroblasts (αSMA), and nuclei (SYTO13) along with a probe atlas designed against 18,676 genes (i.e., the whole human transcriptome). Areas of interest (AOIs) corresponding to specific cell populations within each ROI were separated and the probes extracted by the Bruker’s GeoMx Digital Spatial Profiler (DSP) for library preparation and sequencing. ROIs were classified as immune-hot or immune-cold based on the absolute CD8^+^ T-cell density (CD8/mm² < 100 = cold; ≥100 = hot) (Fig. S1A). We then examined transcriptomic differences between CD8-hot and CD8-cold fibroblast and tumor cell compartments.

**Figure 1.**
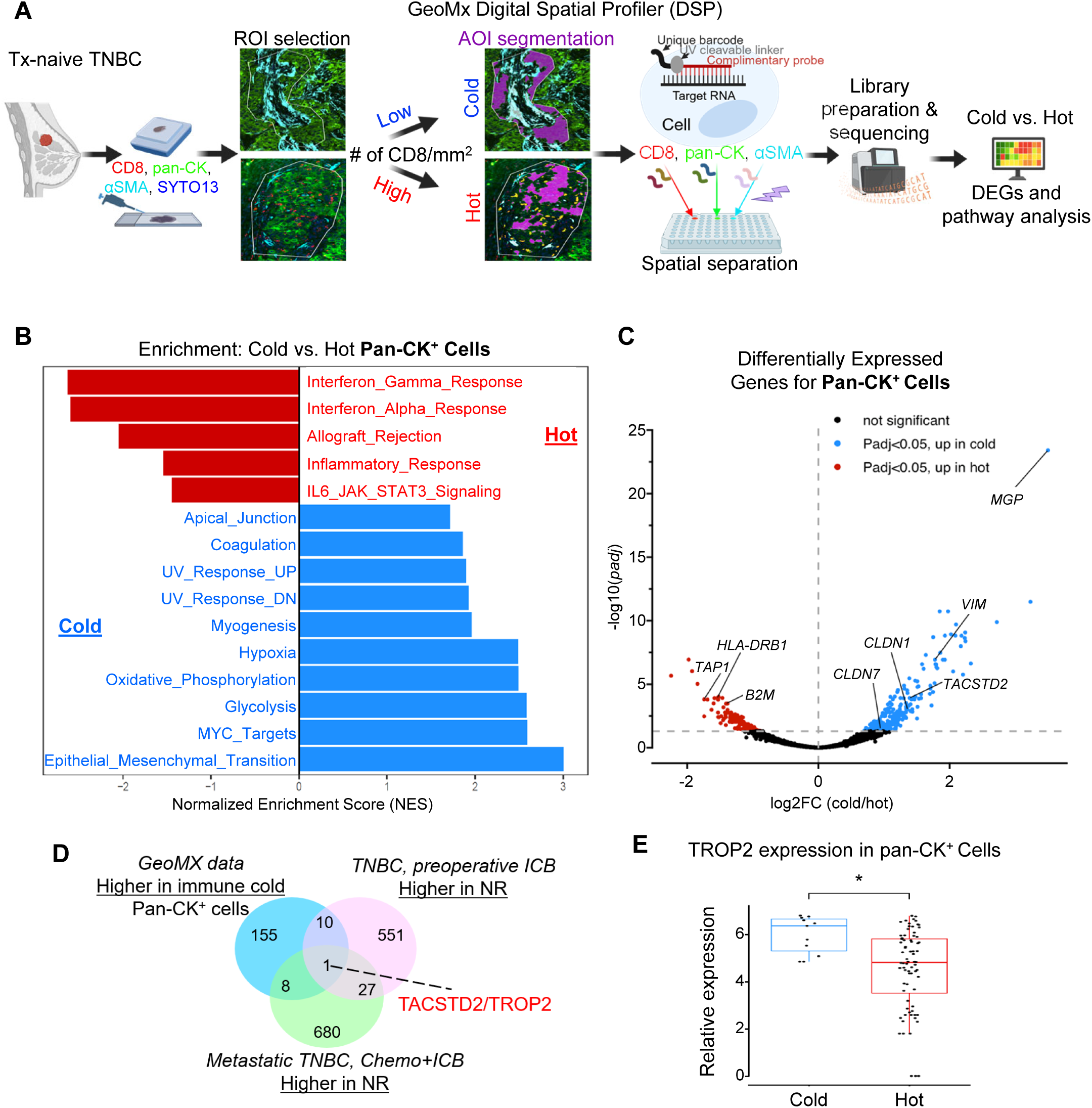
Spatial transcriptomics identifies tumor cell specific programs associated with TNBC immune phenotypes. **A,** Scheme of spatial transcriptomics workflow on the GeoMx Digital Spatial Profiler. Regions of interest (ROIs) were classified as immune-hot or immune-cold based on absolute CD8^+^ T-cell density at the tumor-stroma margin (CD8/mm² < 100 = cold; ≥100 = hot). Within a defined ROI, molecular barcodes are selectively sampled from fluorescence-segmented areas of interest (AOIs) within the instrument. These barcodes are bound to probes targeting 18,675 human RNAs via ultraviolet light-sensitive bonds. Sampling occurs sequentially and non-overlapping, ensuring spatial separation of probes from each AOI. Extracted probes are then used for library preparation and sequencing, generating raw data files that provide RNA transcript counts per gene for each AOI. The data are subsequently analyzed for differential gene expression and correlation studies. **B,** Top 10 pathways associated with pan-CK^+^ cells in the immune cold versus hot ROIs. **C,** Volcano plot of differentially expressed genes between immune cold versus hot pan-CK^+^ cells. **D,** Venn diagram showing genes upregulated in immune-cold pan-CK^+^ cells from GeoMx datasets, genes upregulated in non-responders from a preoperative immune checkpoint blockade (ICB)/pembrolizumab TNBC cohort (30), and genes upregulated in non-responders from a metastatic TNBC cohort treated with chemotherapy plus ICB/pembrolizumab (31). NR: non-responders. **E,** Relative expression level of TACSTD2/TROP2 in the GeoMx dataset.

This analysis revealed that αSMA^+^ fibroblasts in immune-cold tumors were enriched in matrix remodeling and collagen organization programs (Fig. S1B, C), consistent with our recent findings that collagen alignment is a key determinant of immune exclusion (5). In contrast, fibroblasts in CD8-hot tumors exhibited upregulation of antigen presentation and immune-stimulatory pathways (Fig. S1B, C). When analyzing pan-CK^+^ tumor cells, pathways associated with CD8-hot tumors included interferon signaling and allograft rejection (Fig. 1B, C). However, CD8-cold tumors displayed higher expression of epithelial-mesenchymal transition (EMT), myogenesis, and apical junction-related genes including CLDN1, CLDN7, and TACSTD2, encoding TROP2 (Fig. 1B, C).

To prioritize critical genes for further study, we cross-referenced our spatial transcriptomics findings with two clinical TNBC gene expression datasets comparing non-responders versus responders to ICB (30) or chemotherapy plus ICB (31). TROP2 was the only gene consistently shared across all three datasets, highlighting its robustness as a hallmark of immune-cold tumors (Fig. 1D, E).

Given substantial controversy regarding TROP2 function in cancer, we next used an unbiased approach to define pathways associated with TROP2 expression itself across human breast tumors. We employed Gene Set Variation Analysis (GSVA), an adaptation of Gene set enrichment (GSE) that is robust and highly suited to the inter-patient heterogeneity characteristic of human cancer (32). Consistent with our results above, an immunity pathway was the most negatively correlated pathway with TROP2 expression in breast cancer. Notably, the pathway most positively associated with TROP2 was tight junctions (Fig. 2A). We therefore hypothesized that TROP2 expression might play a causal role in influencing the tumor microenvironment in breast cancer. We next interrogated an independent tumor dataset using TIMER, which infers cell populations from bulk RNAseq (33). Again, TROP2 expression was strongly inversely correlated with T cell and immune effector genes including PD1 (PDCD1), granzyme (GZMB) and perforin (PRF1), in multiple breast cancer subtypes, but most strongly in TNBC (Fig. 2B). We also confirmed that TROP2 was negatively associated with cytotoxic T lymphocyte level in a distinct cohort using Tumor Immune Dysfunction and Exclusion (TIDE) analysis (9) (Fig. S2A). Given that lymphocyte counts are prognostic in TNBC (34, 35), we assessed the association of TROP2 levels with relapse-free, metastasis-free and overall survival in basal-like cancers, which are primarily TNBC. High TROP2 expression in each case predicted poor outcome and short survival (Fig. 2C and Supplementary Fig. S2B, C). Thus, these analyses collectively indicate that TROP2 is associated with a paucity of T cells and poor outcomes in triple-negative and basal-like breast cancers.

**Figure 2.**
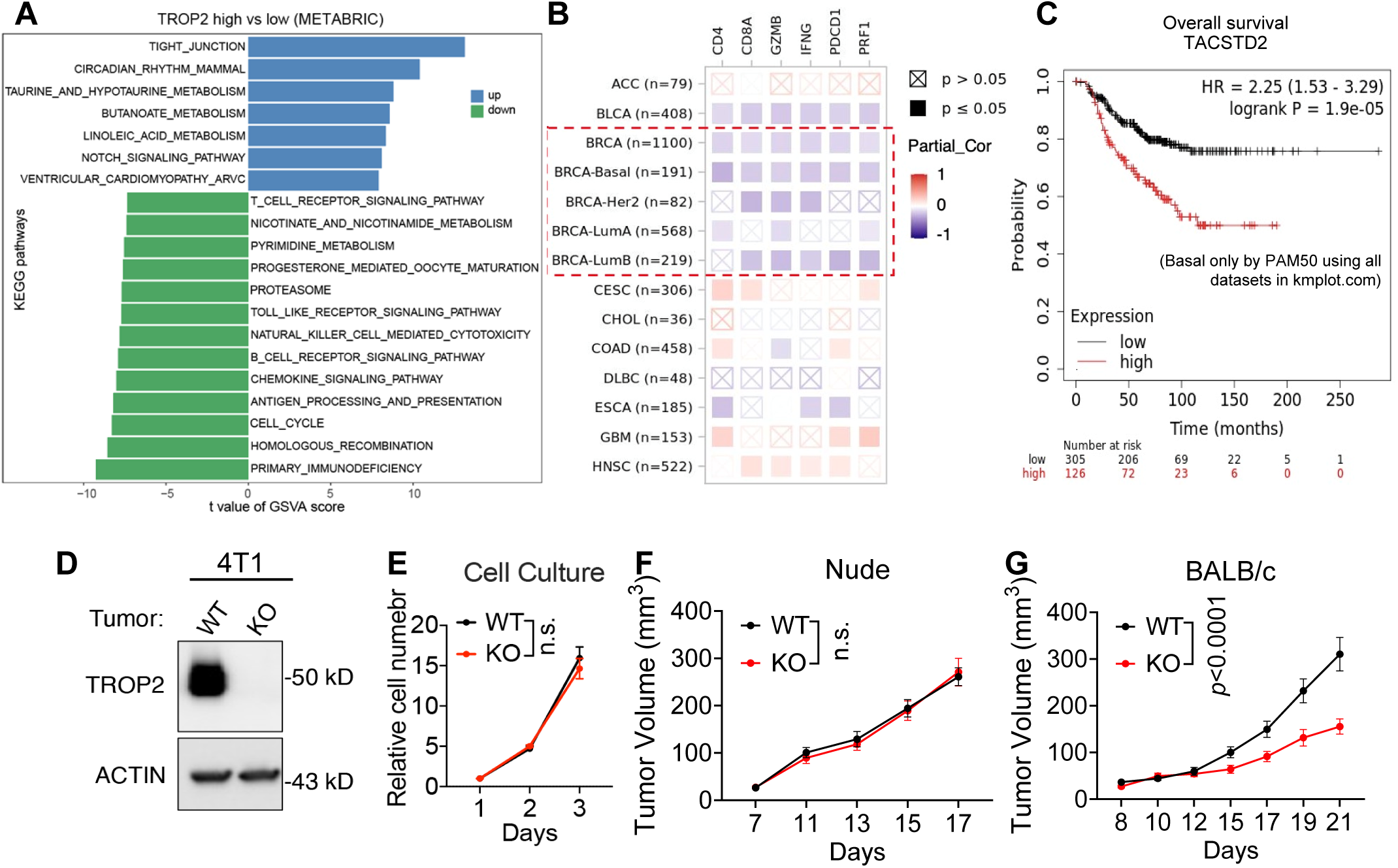
TROP2 promotes mammary tumor growth in immunocompetent hosts. **A,** Top 20 pathways positively (blue) or negatively (green) associated with TROP2 expression in human breast cancer identified by Gene Set Variation Analysis (GSVA) using METABRIC dataset. Top and bottom quartiles of TROP2 expression were used to compute the association. **B,** Correlation between TROP2 and immune cell markers in multiple cancer types by TIMER analysis in the TCGA database. Red dashed box indicates breast cancers. **C,** TROP2 predicts poor overall survival in basal subtype of breast cancer. Generated by Kaplan-Meier Plotter database (https://kmplot.com/analysis/). **D,** Western blot for TROP2 in wildtype (WT) and *Trop2* knockout (KO) 4T1 cells. **E-G,** WT and KO 4T1 tumor cell growth in cell culture (E); in immunocompromised (nude) hosts (n = 10 tumors per group) (F), and in syngeneic immunocompetent BALB/C hosts (WT, n = 12 tumors; KO, n = 10 tumors) (G).

To understand the mechanisms of TROP2 in TNBC we next employed multiple TROP2-expressing syngeneic TNBC models. We ablated endogenous TROP2 via CRISPR-Cas-9 (Fig. 2D), then tested effects in vitro and in vivo through mammary fat pad injection using both immunodeficient and immunocompetent hosts. Consistently, loss of TROP2 had no significant effect on proliferation in vitro, or on tumor growth or apoptosis in immunodeficient nude and NSG mice (Fig. 2E and F, Supplementary Fig. S3A-C). In contrast, TROP2 loss substantially attenuated tumor growth in immunocompetent BALB/c mice compared to control tumors harboring a Cas9/non-targeting gRNA control vector (Fig. 2G). These data suggest that TROP2 promotes tumor growth in an immune-dependent manner.

Immunohistochemistry analysis revealed few CD3^+^, CD4^+^ and CD8^+^ T cells in TROP2-expressing tumors (TROP2-WT), whereas TROP2-knockout (TROP2-KO) tumors were significantly more heavily infiltrated (Fig. 3A and B, Supplementary Fig. S3D-F). Notably, effects of TROP2-KO on tumor growth and lymphocyte infiltration were consistent across both murine TNBC models (Fig. 2-3 and Supplementary Fig. S3G-M). Using flow cytometry, we analyzed T cell subpopulations in TROP2-WT and TROP2-KO tumors. TROP2-KO tumors exhibited significant increases in effector memory CD8^+^ and CD3^+^ T cells (CD44^+^/CD62L^-^), and PD1^+^ CD8^+^ and CD3^+^ T cells indicative of activation rather than exhaustion, as well as Granzyme^+^ CD8^+^ cells (Fig. 3C-H and Supplementary Fig. S3L and M). Consistent with these findings, immunohistochemistry also showed an increase in the T cell activation marker 4-1BB^+^ and CD69^+^ cells in KO tumors compared with WT tumors (Supplementary Fig. S3N-Q). Furthermore, RNAseq analysis of the bulk tumors deconvoluted using CIBERSORT and TIMER consistently demonstrated significant increases in CD8^+^ T cells (Supplementary Fig. S4A). Collectively, these data show that loss of TROP2 results in infiltration of effector memory and activated T cells in association with attenuated breast tumor progression.

**Figure 3.**
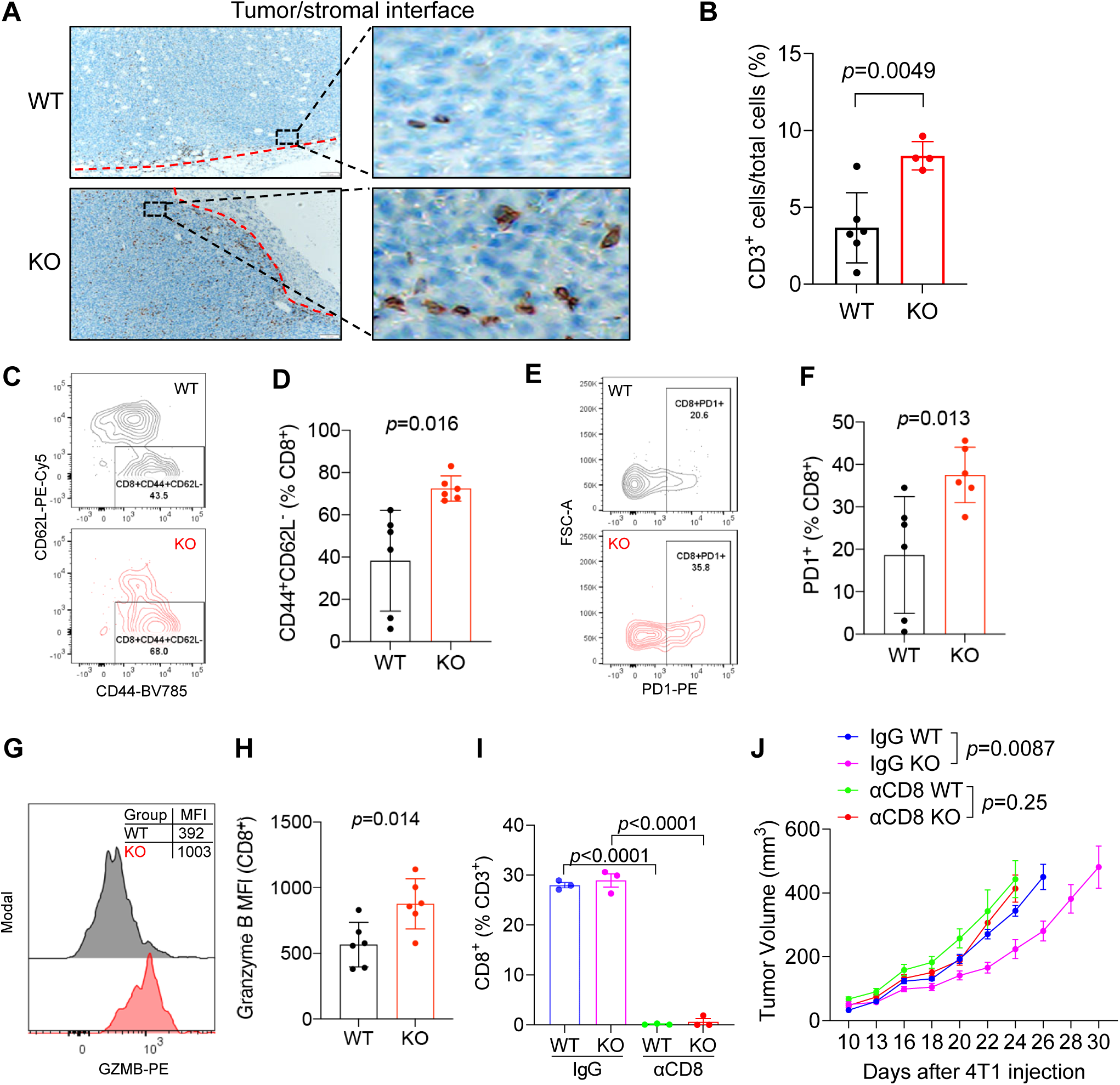
TROP2 promotes mammary tumor immune exclusion. **A** and **B,** Representative images (A) and quantification (B) of CD3^+^ T cell IHC staining in WT and KO tumors. **C** and **D**, Representative flow cytometry contour plots (C) and quantification (D) of CD44^+^ CD62L^-^ effector memory CD8^+^ T cells in WT and KO tumors. **E** and **F**, Representative flow cytometry contour plots (E) and quantification (F) of PD1^+^ activated CD8^+^ T cells in WT and KO tumors. **G** and **H**, Representative flow cytometry histogram (G) and quantification (H) of Granzyme B mean fluorescence intensity (MFI) in CD8^+^ T cells in WT and KO tumors. **I,** Percentages of CD8^+^ cells among CD3^+^ T cells in splenocytes harvested from mice treated with control IgG or anti-CD8 depleting antibodies. **J,** *Trop2*-WT and -KO 4T1 tumor growth curves in BALB/C mice treated with control IgG or anti-CD8 depleting antibodies (WT, n = 10 tumors per group; KO/IgG, n = 8 tumors; KO/anti-CD8: n = 6 tumors).

We then tested directly the contribution of cytotoxic T cells in the setting of TROP2 loss. We carried out IgG-mediated CD8^+^ T cell depletion in mice bearing TROP2-WT or matched TROP2-KO TNBC tumors (Fig. 3I). In mice treated with control IgG, tumors with loss of TROP2 displayed a substantial growth disadvantage as we had observed previously. However, in the setting of CD8 depletion the difference in growth between TROP2-KO and TROP2-WT tumors was abolished (Fig. 3J). Taken together, these findings support a role for TROP2 in immune cell exclusion of CD8^+^ effector cells that drives tumor progression.

We next sought to uncover how TROP2 may instigate immune exclusion and to probe the concordance in TROP2-associated pathways between our murine models and human tumors. Thus, we carried out bulk RNAseq analysis of matched TROP2-WT and TROP2-KO tumors and identified altered transcriptional programs via Gene Set Enrichment Analysis (GSEA). Indeed, we observed agreement in signatures associated with TROP2 expression in human and mouse tumors, as the top gene sets positively enriched in TROP2-WT versus TROP2-KO tumors involved cell-cell contact and tight junctions, whereas the most negatively enriched pathways were those involving inflammation, tumor immune responses, and T cell cytotoxicity (Fig. 4A and Supplementary Fig. S4B). The potential association of TROP2 in this context with tight junctions was particularly notable, as apical/tight junction appeared as one of the top pathways enriched in our human TNBC GeoMx dataset for immune cold tumor cells (Fig 1B).

**Figure 4.**
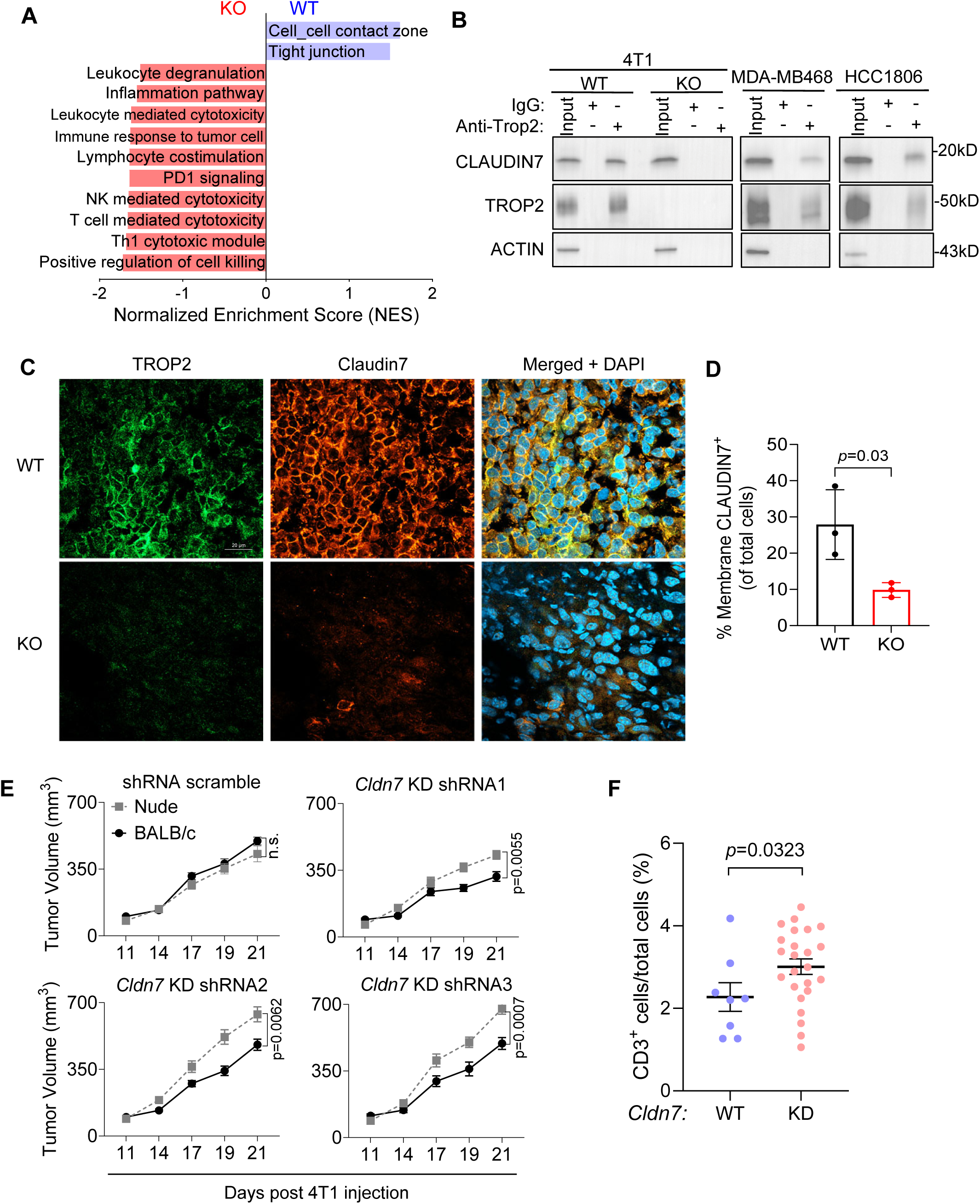
TROP2 promotes a tight junction-mediated barrier and inhibits a pro-inflammatory tumor immune microenvironment. **A,** Enriched pathways from Gene set enrichment analysis (GSEA) on bulk RNA-seq of *Trop2*-WT and KO 4T1 tumors harvested from BALB/C hosts. **B,** Western blot analysis showing co-immunoprecipitation (co-IP) of Claudin7 and TROP2 using IgG or anti-TROP2 in *Trop2*-WT and KO 4T1 (left panel), MDA-MB468 (middle panel), and HCC1806 (right panel) TNBC cells. **C,** Representative tumor tissue immunofluorescence images of TROP2, tight junction molecule Claudin7, and DAPI (nuclear stain), showing TROP2/Claudin 7 co-localization, and loss of Claudin7 expression and membrane localization with TROP2 knockout. **D,** Quantification of proportion of membrane Claudin7-expressing cells in WT and KO 4T1 tumors for panel C. **E,** *Cldn7*-WT (scramble) and - KD 4T1 tumor growth curves in immunodeficient nude or immunocompetent BALB/C mice (n=8 tumors per group). **F,** Quantification of CD3^+^ T cell IHC staining of *Cldn7*-WT and -KD 4T1 tumors harvested from BALB/c mice.

TROP2 is reported to have a highly tissue-selective role in tight junction formation within the normal corneal epithelium, which is disrupted by germline loss-of-function *TACSTD2* mutations (24). In the cornea, TROP2 interacts with Claudin 7, a key tight junction protein whose downregulation is known to result in disruption of these structures (26). Thus, we next tested the physical interaction of TROP2 and Claudin 7, and observed strong co-immunoprecipitation (co-IP) of the two proteins in both human and murine TNBC cells (Fig. 4B and Supplementary Fig. S4C). We then performed immunofluorescence for Claudin 7 in the TROP2-WT and TROP2-KO TNBC tumors. As anticipated, in tumors expressing endogenous TROP2, Claudin 7 showed primarily membrane localization and strong co-localization with TROP2, indicative of intact tight junctions. However, tumors with loss of TROP2 demonstrated substantially decreased expression and abrogated membrane localization of Claudin 7 protein, as well as decreased expression of both Claudin 7 and Claudin 1 mRNA (Fig. 4C, D and Supplementary Fig. S4D). We further confirmed tight junction integrity through analysis of an additional tight junction protein, Occludin (36). While TROP2-WT tumors demonstrated strong membrane Occludin staining, TROP2-KO tumors showed dramatic loss of Occludin staining and localization, as well as decreased mRNA expression (Supplementary Fig. S4D-F). Notably, Claudin 1, 7 and Occludin expression are all highly positively correlated with TROP2 expression in human breast tumors (Supplementary Fig. S4G). Thus, TROP2 physically interacts with Claudin 7 in TNBC, and its loss disrupts tight junctions via altered expression and localization of multiple integral tight junction proteins.

To directly test the contribution of Claudin7 and tight junctions in antagonizing anti-tumor immunity, we generated 4T1 TNBC lines with shRNA-mediated *Cldn7* or control knock down (KD) (Supplementary Fig. S5A), then injected them into immunodeficient nude mice and immunocompetent BALB/c mice. We found that control tumors grew comparably within nude mice and BALB/c mice, while all three *Cldn7*-KD tumor lines grew significantly slower in the BALB/c mice compared to nude mice (Fig. 4E). The *Cldn7*-KD tumors also showed significantly higher CD3 infiltration compared to the control tumors (Fig. 4F and Supplementary Fig. 5B). Thus, disruption of the essential tight junction factor Claudin 7 phenocopies TROP2 loss, impeding in vivo tumor progression in an immune-dependent manner.

We next employed reconstitution experiments to establish whether restoration of TROP2 expression was sufficient to mediate immune exclusion and tumor progression. Prior studies have linked TROP2 signaling and function in cancer to its intracellular domain (ICD), whereas binding to Claudin 7 may be mediated through its transmembrane region (37). Therefore, we tested whether the ICD was required to promote immune exclusion and tumor progression by generating a truncated TROP2 protein containing the extracellular and transmembrane domains (TROP2-ECDTM) but lacking the ICD. We reconstituted TROP2-KO TNBC cells with either full-length TROP2 or the TROP2-ECDTM truncation mutant, and confirmed by flow cytometry that both proteins were localized to the cell surface (Fig. 5A). We found that indeed the truncated TROP2-ECDTM physically associated with Claudin 7 by co-IP (Supplementary Fig. S5C, D). Neither full-length TROP2 nor the ECDTM mutant affected tumor cell proliferation compared to TROP-2 KO cells in vitro (Fig. 5B). We then implanted the TROP2-KO or the TROP2 reconstituted cells into syngeneic hosts. TROP2 reconstitution strongly promoted tumor growth in vivo, and most importantly reconstitution with TROP2-ECDTM conferred a similar growth advantage (Fig. 5C). Furthermore, TROP2-ECDTM reconstitution fully restored Claudin 7 expression to the level seen in full length TROP2-reconstituted tumors (Fig. 5D and E), and this effect was associated with a substantial decrease in T cell infiltration in reconstituted tumors (Fig. 5F and G). These findings demonstrate that the C-terminal intracellular domain of TROP2 is dispensable for Claudin 7 interaction and expression that are associated with immune exclusion and TNBC progression in vivo.

**Figure 5.**
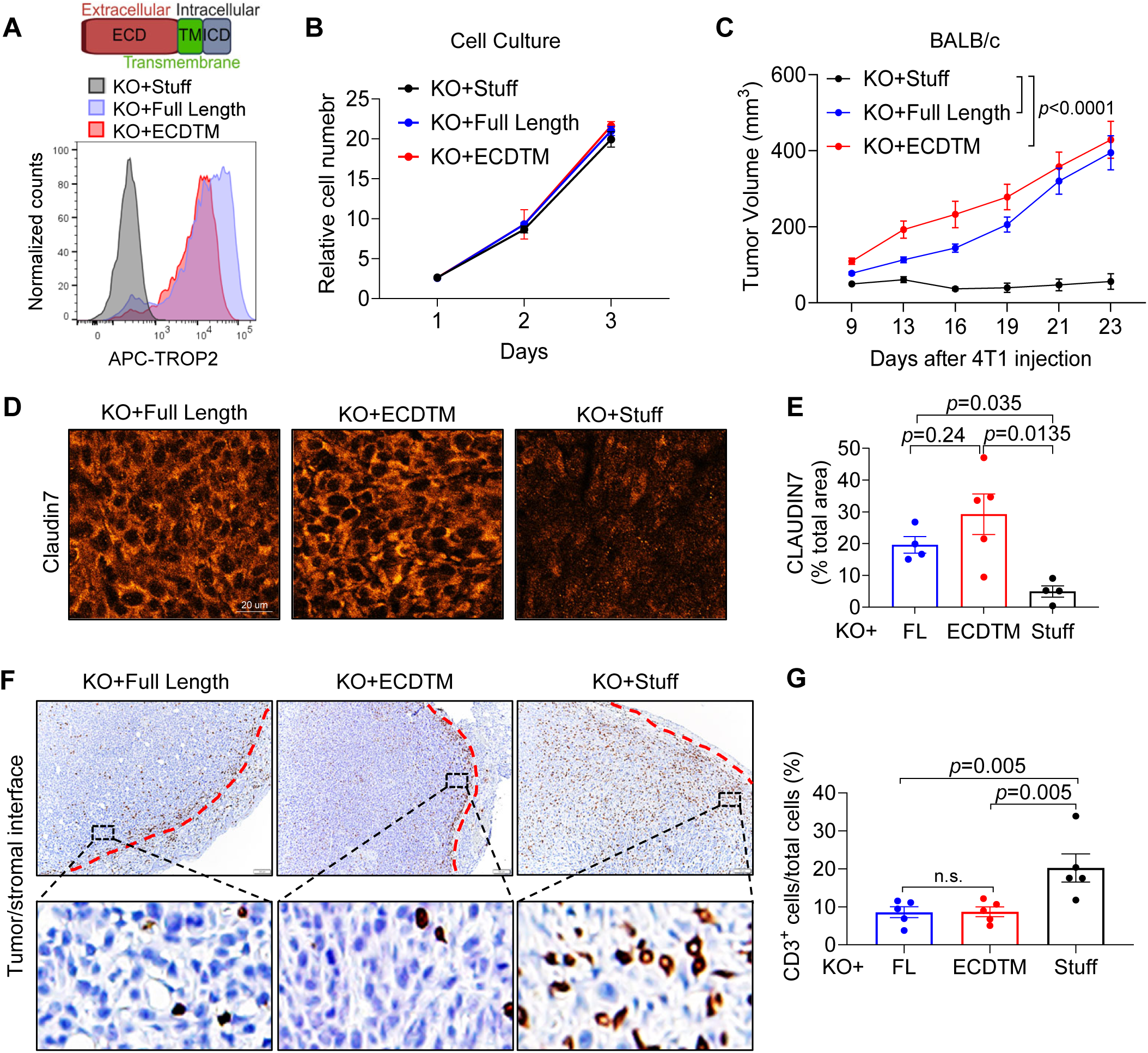
TROP2 intracellular domain is dispensable for its tight junction and immune barrier function. **A,** Histogram showing flow cytometry analysis of cell-surface TROP2 in *Trop2*-KO 4T1 reconstituted with control vector (Stuff), Full-Length TROP2, or C-terminal truncated ECDTM mutant. **B,** In vitro proliferation of *Trop2*-KO 4T1 reconstituted with control vector, Full-Length TROP2 or ECDTM truncated mutant. **C,** Tumor growth curves of *Trop2*-KO 4T1 reconstituted with control vector (Stuff), Full-Length TROP2, or C-terminal truncated ECDTM TROP2 mutant in BALB/C immunocompetent hosts (n = 10 tumors per group). **D,** Representative tumor tissue immunofluorescence images of Claudin7 in tumors harvested from panel C. **E,** Quantification of Claudin7 for panel D. **F** and **G**, Representative images (F) and quantification (G) of CD3^+^ T cell IHC staining in tumors harvested from panel C.

Based on these findings, we sought to therapeutically exploit the role of TROP2 in immune exclusion by testing effects of TROP2 targeting on response to PD1 checkpoint blockade. Sacituzumab govitecan (SG) is a TROP2 directed antibody drug conjugate (ADC) comprised of the hRS7 antibody component, whose binding to TROP2 on the cell surface results in its rapid internalization, linked to a topoisomerase 1 inhibitor (38). To circumvent complications induced by the SG cytotoxic payload topoisomerase 1 inhibitor, we choose to employ naked hRS7 antibody in combination with PD1 checkpoint blockade. This experiment required reconstitution of TROP2-KO TNBC cells with human TROP2 (hTROP2), as hRS7 does not interact effectively with the murine protein. We first demonstrated that hTROP2 interacts strongly by co-IP with murine Claudin 7 in these cells (Fig. 6A and B). Accordingly, following implantation of cells reconstituted with either control vector or hTROP2 into syngeneic hosts, we observed a significant growth advantage of the hTROP2-expressing tumors in immunocompetent mice (Supplementary Fig. S6A). Furthermore, analogous to the effects of murine TROP2 reconstitution, while TROP2-KO tumors reconstituted with control vector were heavily infiltrated with T cells, hTROP2-reconstitued tumors demonstrated a paucity of T cells (Supplementary Fig. S6B and C). We then generated larger cohorts of tumor-bearing mice and carried out a 4-arm experiment, treating with control vehicle, hRS7, anti-PD1, or the combination. Single-agent hRS7 did not significantly affect tumor progression, indicating little or no cytotoxic effect of hRS7 alone at doses used. Similarly, anti-PD1 alone had little effect in this relatively poorly immunogenic model. In contrast, the combination of hRS7 and anti-PD1 significantly impeded tumor progression in hTROP2-reconstituted tumors (Fig. 6C). Importantly, immunofluorescence staining and immunohistochemistry showed a significant reduction of Claudin7 and increased CD3^+^ cells, respectively, in the hRS7-treated tumors compared to vehicle group (Fig. 6D-G), suggesting a disrupted tight junction barrier and increased permeability of immune infiltration. In addition, the anti-PD1 and combination treatment significantly enhanced the proportion of early activated TIM3^-^PD1^+^CD8^+^ T cells (Fig. 6H, I). To test the causal role of Claudin7 in this phenotype, we engineered doxycycline inducible *Claudin7* knockdown 4T1 cells to reduce Claudin7 expression in established tumors (Supplementary Fig. S7A), thus mimicking the hRS7 mediated effect. We found that upon doxycycline treatment, 4T1 tumors become sensitive to an anti-PD1 treatment, associated with significantly increased TIM3^-^PD1^+^CD8^+^ T cells (Supplementary Fig. S7B-C). Thus, TROP2 targeting and immune checkpoint blockade function through different mechanisms involving increased T cell infiltration and activation, respectively. Collectively, these data demonstrate that neither recruitment nor activation alone is sufficient to induce an anti-tumor effect. Instead, the data reveal that only by increasing both the number of recruited T cells (by blocking TROP2) and their activation state (by blocking PD1) is an optimal response achieved.

**Figure 6.**
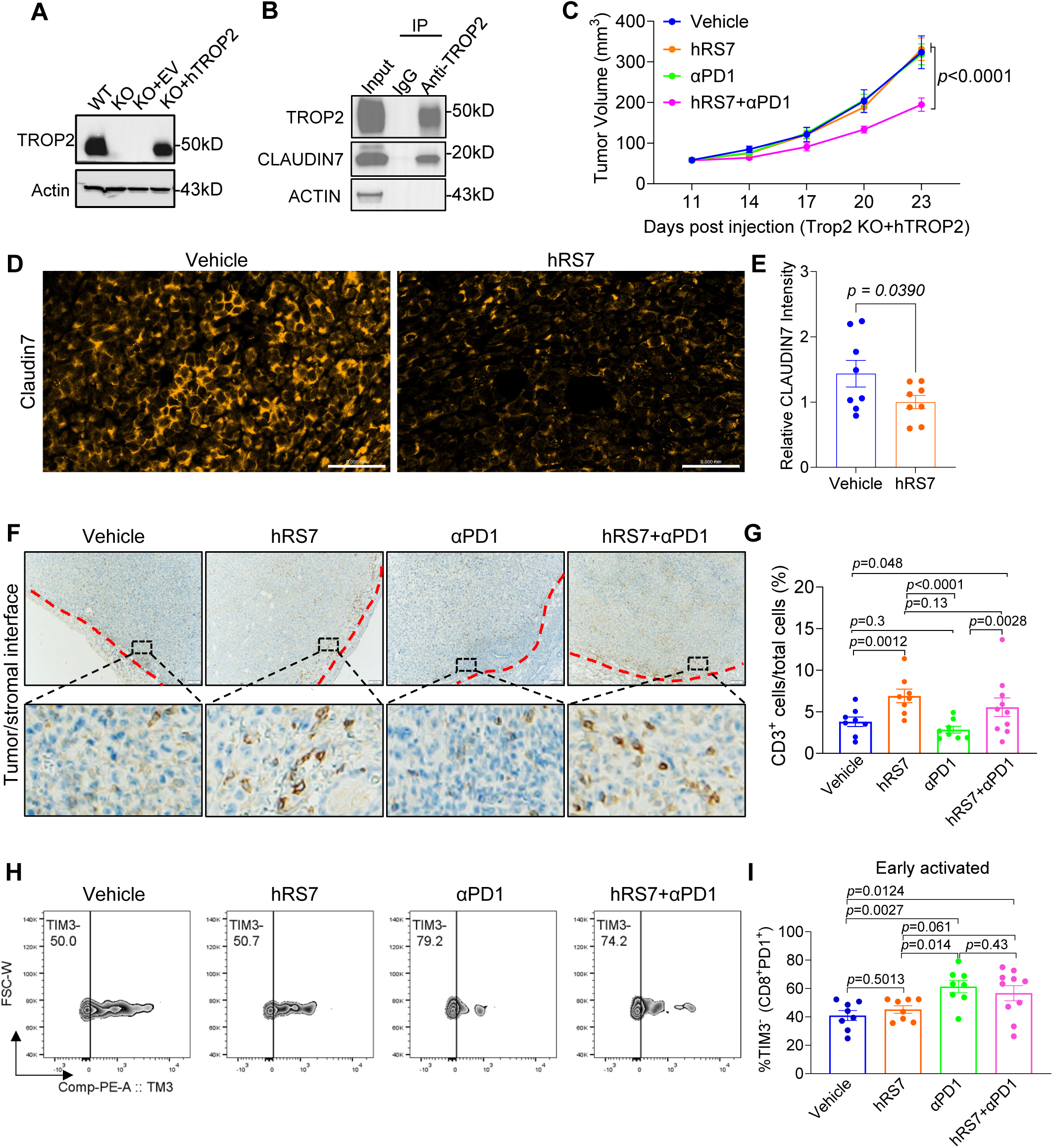
TROP2 targeting via the anti-TROP2 antibody component of SG promotes anti-PD1 efficacy. **A,** Western blot of *Trop2*-KO 4T1 cells reconstituted with empty vector (EV) or human TROP2 (hTROP2). **B,** Western blot showing co-IP of hTROP2 and Claudin 7 in *Trop2*-KO 4T1 cells reconstituted with hTROP2. **C,** Tumor growth curves for KO+hTROP2 (hRS7+anti-PD1, n = 10 tumors; n = 8 tumors per group for other three groups) in BALB/C mice treated with Vehicle, hRS7, anti-PD1, or hRS7+anti-PD1 combination. Treatment started when tumors reached 100 mm^3^. Arrows indicate treatment days. **D,** Representative tumor tissue immunofluorescence images of Claudin7 in tumors harvested from panel C. **E,** Quantification of Claudin7 for panel D. **F** and **G**, Representative images (F) and quantification (G) of CD3^+^ T cell IHC staining in tumors harvested from panel C. **H** and **I,** Representative flow cytometry contour plots (F) and quantification (G) of TIM3^-^ early activated CD8^+^ PD1^+^ T cells in tumors harvested from panel C. \**p*<0.05, \*\**p*<0.01.

Lastly, we investigated the contribution of TROP2 to immunotherapy response in human breast cancer. We identified cohorts with available pre-treatment tumor RNAseq data in which breast cancer patients were treated with anti-PD1 therapy (Pembrolizumab), either with or without chemotherapy (Cohorts 1 and 2, respectively) (30, 31). We examined the impact of TROP2 expression on treatment effect through bulk/pseudo-bulk RNA analysis, plotting the odds ratio (OR) for effect as a function of TROP2. We found that for both cohorts high TROP2 expression was significantly negatively associated with treatment effect (OR for Pembrolizumab plus chemotherapy Cohort 1: 0.06, 95% CI 0.0-0.61; OR for Pembrolizumab alone Cohort 2: 0.06, 95% CI 0.01-0.36) (Fig. 7A). In these same datasets, expression of the highly homologous epithelial protein EpCAM was not significantly associated with treatment effect in either cohort, and PD-1 expression was significantly positively associated with effect more strongly in Cohort 2 than Cohort 1 (Fig. 7A). We also took advantage of Cohort 2 single-cell RNAseq (scRNAseq) data to demonstrate that TROP2 expression specifically among epithelial cells was negatively associated with treatment effect (OR: 0.15, 95% CI 0.02-0.84), while again EpCAM was not (Supplementary Fig. S8A-B). Furthermore, a significant overlap and positive correlation between TROP2 and CLDN7 expression was observed selectively in the non-responsive group (Supplementary Fig. S8C). Finally, when we analyzed the subset of cells identified as CD8 T cells and stratified the patient tumors by epithelial TROP2 expression, we found that CD8 T cells from tumors with high TROP2 exhibit primarily a naïve signature, while those from tumors with low TROP2 had higher interferon response and cytotoxicity signatures (Fig. 7B). These results collectively support our conclusion that TROP2 expression promotes a program of T cell exclusion, resulting in diminished immune cell engagement and persistence in a naïve state, while low TROP2 tumors are more accessible to immune cells, resulting in activation and cytotoxicity that mediate anti-tumor immunity (Fig. 7C).

**Figure 7.**
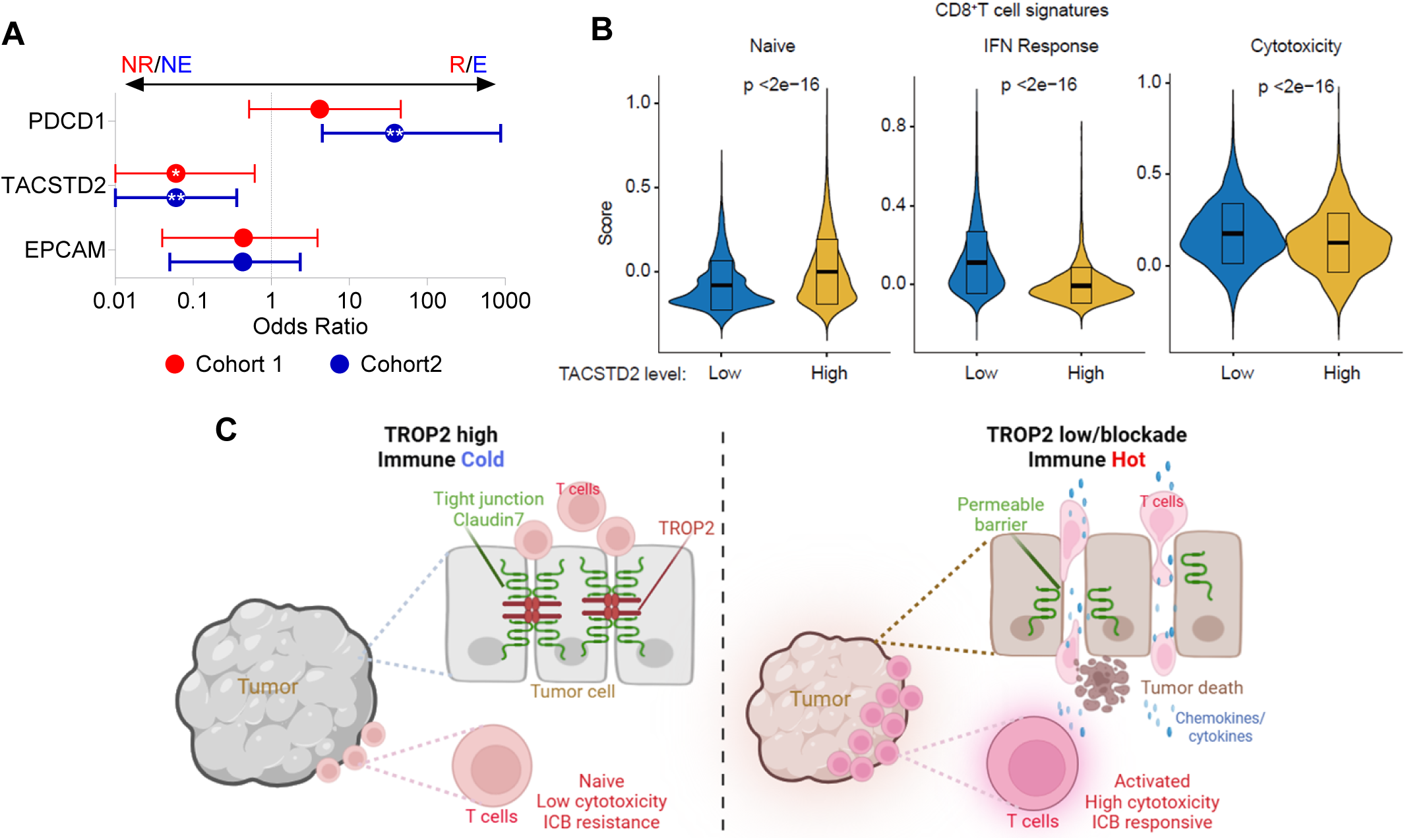
High TROP2 associates with poor response to immune checkpoint inhibitors in human breast cancer. **A,** Odds ratios for associations between gene expression (PDCD1 (PD1), TACSTD2 (TROP2) and EPMCAM) and response to anti-PD1 therapy. Cohort 1: Response (evaluated radiologically using RECIST criteria) to combination Pembrolizumab, Carboplatin, and Nab-paclitaxel among patients with advanced TNBC (31); NR: non-responsive, R: responsive. Cohort 2: Response assessed by T cell receptor clonotype expansion following pre-operative single-agent Pembrolizumab for breast cancer (30), NE: no clonotype expansion, E: yes clonotype expansion. **B,** Violin plots derived from single-cell RNAseq data from the cohort in (A), showing subset signature scores among CD8^+^ T cells in tumors expressing low vs. high epithelial TROP2 (30). CD8 signatures are derived from PMID: 37248301. **C,** Proposed model for TROP2-mediated immune exclusion in breast cancer. Left, TROP2 high enables a claudin/tight junction-mediated mechanical barrier to exclude antitumor immune infiltration. Right, TROP2 low or pharmacological blockade deregulates the tight junction-mediated mechanical barrier and allows antitumor immune infiltration. ICB: immune checkpoint blockade.

## DISCUSSION

Our study reveals a heretofore unidentified role of TROP2 expression in cancer, enabling a tight junction-based barrier mechanism to exclude T cells and evade anti-tumor immunity. Our conclusions are supported by analysis of human tumors, by gain- and loss-of-function studies of TROP2, by comparison of immunodeficient and immunocompetent syngeneic models, and by direct T lymphocyte depletion experiments. Further supporting the link between TROP2 and tight junctions in promoting tumor progression, we find that Claudin 7 depletion in TNBC mimics loss of TROP2 in its immune-specific effects in vivo. These findings have implications beyond TROP2 itself, as they underscore a potentially broad role for tight junctions as a mechanism of barrier-mediated immune exclusion in human cancer. Accordingly, this work suggests the possibility that targeting not only TROP2 but also other tight junction-associated proteins in the proper context could be fruitfully employed to reverse immune evasion and enhance anti-tumor immune responses. Although our study focuses on breast cancer, TROP2 may play a similar role in control of tight junctions, the immune microenvironment and checkpoint blockade response in other tumor types that exhibit aberrant TROP2 expression, including melanoma, ovarian and lung cancers (3, 39).

The mechanisms governing immune exclusion and evasion are multifaceted, often varying significantly across tumor types and even within specific subtypes. For instance, HR^+^ breast tumors generally exhibit greater genomic stability and a lower tumor mutational burden compared to TNBC, resulting in reduced immunogenicity and a less infiltrated microenvironment (40). While most epithelial tumors, including TNBC, lack the organized architecture of normal epithelia, they frequently exploit specific mechanisms of immune exclusion. In various malignancies, cancer-associated fibroblasts drive a collagen-rich stromal reaction that physically blocks T-cell entry (41). Our findings identify TROP2 expression as a novel contributor to barrier-mediated immune exclusion in TNBC. Although TROP2 expression may decrease in tumors undergoing epithelial-mesenchymal transition (EMT), such tumors often employ alternative exclusion strategies. For example, EMT is closely linked with TGF-beta signaling, which can directly inhibit T-cell migration and stimulate CAFs to remodel the extracellular matrix. Thus, TROP2-mediated exclusion represents a distinct regulatory layer within the complex architecture of the TNBC immune landscape.

Intriguingly, we do not observe significant cell-autonomous effects of TROP2 knockout on TNBC cell proliferation in vitro or tumor progression in immunodeficient models. Furthermore, we find that reconstitution of knockout cells with a truncated TROP2 mutant lacking the intracellular signaling domain is sufficient to fully restore claudin integrity and tumor growth in vivo. While our findings do not exclude a potential contribution of the TROP2 intracellular domain and its signaling in some contexts, these results highlight for the first time a central role of claudin interaction and tight junction-mediated immune exclusion as drivers of tumor progression mediated by TROP2 (15).

The potential ability of TROP2 to control tight junction integrity and immune exclusion in multiple cancer types contrasts with the highly tissue-specific corneal tight junction dysregulation associated with germline TROP2 mutation (26). While the precise mechanism of this tissue specificity is not clear, it could possibly be explained by functional redundancy between TROP2 and its paralogue EpCAM for claudin regulation in non-tumor contexts (42, 43). In TNBC, we find that loss of TROP2 is sufficient to disrupt claudin expression and localization in vivo, and that claudin is restored by TROP2 reconstitution. Further underscoring the central role of TROP2 for tight junction regulation in TNBC, we find that TROP2-KO tumors also display significantly lower Claudin 1/7 and Occludin mRNA than TROP2-WT tumors, potentially mediated by a recently-described positive feedback loop between tight junction barrier dysfunction and inflammation (44). In contrast to TROP2, multiple lines of evidence argue against a major role for EpCAM in driving immune exclusion in TNBC. First, unlike TROP2 and Claudin 7, EpCAM was not among the top genes enriched in immune-cold versus immune-hot pan-CK^+^ cells (Fig. 1C, D). Second, TIMER analysis revealed no strong inverse association between EpCAM expression and T cell markers in breast cancer (Supplementary Fig. S8D), contrasting with the clear relationship observed for TROP2 (Fig. 2B). Third, as shown in Fig. 7A, EpCAM expression did not correlate with response to immune checkpoint blockade in breast cancer, whereas TROP2 demonstrated a statistically significant association.

Despite a heretofore limited understanding of its tumor-specific functions, TROP2 has emerged as an important target for anti-cancer ADC therapy. The TROP2-targeted ADC SG has improved overall survival for both advanced TNBC (29) and hormone-receptor positive breast cancer (45). Nonetheless, the contribution to therapeutic response of TROP2 targeting per se, versus delivery of cytotoxic payload have not been extensively explored. We show directly that TROP2 impedes tumor infiltration by immune cells in TNBC models, and that loss of TROP2 enables recruitment of key effector memory, PD1^+^ and activated CD8^+^ T cell subsets. Correspondingly, we find that blocking TROP2 with the TROP2-directed antibody component of SG sensitizes refractory TNBC tumors to PD1 inhibition in vivo. In addition, this dual TROP2/PD1 blockade promotes both intra-tumoral accessibility and activation of antitumor T cells. Supporting the clinical relevance of these findings, we demonstrate that TROP2 levels are a negative predictor of treatment response as assessed by tumor regression and clonal T cell expansion for breast cancer patients receiving anti-PD1 therapy, including single-agent Pembrolizumab without chemotherapy (30, 31).

The role defined herein for TROP2 in tight junction-mediated immune exclusion has broad implications for how TROP2-targeted and other ADCs are employed in the future. Multiple tumor types appear to exhibit TROP2-associated immune exclusion (Fig. 2B), and these cancers would be reasonable candidates for therapeutic testing with emerging ADCs targeting not only TROP2 but also other tight junction components, potentially in combination with immune checkpoint blockade, as exemplified by the recent successful phase 3 clinical trial demonstrating superiority of sacituzumab govitecan plus pembrolizumab compared to standard chemotherapy plus pembrolizumab for TNBC (46). An immune-mediated effect linked to TROP2/tight junction targeting also highlights the potential for the next generation of ADC designs. These could include the possible incorporation of bispecific immune cell engagers, and immune-activating rather than cytotoxic payloads. Finally, the pivotal link exemplified by TROP2 between intercellular junctions as mechanical barriers and anti-tumor immune exclusion opens up additional new opportunities for innovative investigation and therapeutic development in the field of anti-cancer immunity.

## METHODS

### Spatial transcriptomics data generation

Archived, paraffin-embedded, treatment-naïve TNBC biopsies from eight patients were used as the primary material for this study. Serial sections were prepared from these blocks, with the first section stained with hematoxylin and eosin (H&E) and reviewed by two pathologists (V.I.B. and B.K.P.) for morphological assessment. The second section was stained with probes against 18,676 human RNAs, covering the entire transcriptome (NanoString Technologies, 121401102) and multiplex immunofluorescence staining to identify cell types of interest. These fluorescent morphology markers included the nuclear stain SYTO13 (AF488 conjugated, NanoString Technologies), PanCK for tumor epithelium (AF532 conjugated, NanoString Technologies, 121300310), CD8a for CD8^+^ T lymphocytes (AF594 conjugated, Clone: C8/144B; Biolegend, 372904) and αSMA for fibroblasts (AF647 conjugated, Clone: 1A4; R&D Biologicals, IC1420R-100UG). H&E morphology of the first slide was used to visually assess for varied immune density and optimum tumor epithelium on the GeoMx Digital Spatial Profiler instrument (NanoString Technologies) to mark suitable regions of interest (ROIs) on the probe- stained slide. These ROIs were further segmented by the instrument because of differing fluorescence into distinct areas of interest (AOIs) corresponding to tumor, immune and fibroblast populations. Segmentation accuracy was verified before finalizing barcode collection. Barcodes, linked to oligonucleotides via ultraviolet light-cleavable bonds, were captured in designated wells of a 96-well collection plate. Spatial transcriptomic mapping was achieved by sequential collection and well-based separation of AOIs. The unique barcodes were subsequently subjected to Illumina-based sequencing following library preparation on their NextSeq1000 platform with a 100 cycle P2v3 flow cell. Target probe-specific count files per AOI were generated and downstream bioinformatic analyses undertaken with the R Computing platform using appropriate software packages.

### Computational analysis

NanoString GeoMx NGS data was processed following standard guidelines using the GeomxTools R package (47). AOIs detecting <10% of genes and genes detected in <10% of AOIs were excluded, resulting in 155 of 190 AOIs and 14,833 of 18,676 genes retained for analysis. Differential expression analysis was performed using the limma R package (48). First, effective library sizes for each AOI were estimated using the trimmed mean of M-values (TMM) method via the calcNormFactors function from edgeR. The count data was then normalized using limma’s voom function. Normalized gene expression was modeled with infiltration identity (cold vs. hot) and patient identity as fixed predictor variables. Differential expressions were assessed for the following linear contrasts: cold vs. hot tumor AOIs and cold vs. hot fibroblast AOIs. For each comparison, gene-level significance was determined using Benjamini-Hochberg correction, the default in limma.

For deconvolution and abundance analysis, counts data were quantile (Q3)-normalized and analyzed using the SpatialDecon R package. A custom reference profile was derived from breast cancer scRNA-seq data (49), incorporating the top differentially expressed fibroblast marker genes (FDR < 0.05). The spatialdecon function estimated fibroblast sub-type proportions in each AOI. A Poisson regression model (glm function) was used to assess cell-type count differences between immune-hot and immune-cold AOIs, with statistical significance determined using a Local Wald Test, implemented with the multcomp R package. All p values were corrected with Bonferroni adjustment.

Gene set enrichment analysis (GSEA) was performed using the clusterProfiler R package (50). For each of the two linear contrasts tested during differential expression analysis, the obtained results were pre-sorted by t statistic value. GSEA was then implemented with the fgsea algorithm on this pre-sorted gene list for the following gene set collections: Hallmark gene sets from MSigDB and Biological Processes gene sets from Gene Ontology. Significance was assessed after all p values were corrected with Benjamini-Hochberg adjustment.

### Cell Lines

Trop2 was knocked out in mouse mammary tumor cell lines 4T1 (MGH Center for Molecular Therapeutics Cell Bank) and AT-3 (Sigma-Aldrich, SCC178) using TROP2 sgRNA CRISPR/Cas9 All-in-One Lentivector set (ABM, 460591140595). A CRISPR/Cas9 vector with non-targeting sgRNA (ABM, K010) was used to generate control *Trop2*-WT cells. Cldn7 was knocked down using following shRNA sequences in the pLV[shRNA]-Bsd-U6 lentiviral backbone: shRNA1: GACGCCCATGAACGTTAAGTA; shRNA2: CATCAGATTGTCACAGACTTT; shRNA3: AGCGAAGAAGGCCCGAATA. shRNA scramble: CCTAAGGTTAAGTCGCCCTCG was used as the control. In short, lentiviral packaging was performed by cotransfecting HEK293T cells with CRISPR/Cas9 vector and LV-MAX Lentiviral Packaging Mix (Gibco, A43237V) using Lipofectamine2000 (Invitrogen, 11668-019). After 48h, lentivirus-containing medium was collected, filtered, and used to infect the target tumor cell lines. The infected cells were then expanded and selected by puromycin. Human triple negative breast cancer cell lines MDA-MB468 and HCC1806 were obtained from the MGH Center for Molecular Therapeutics Cell Bank.

The mouse TROP2 cDNA was used to generate expressing plasmids (VectorBuilder) carrying full length TROP2 and TROP2 ECDTM truncated version. Control vector carrying same lentiviral elements but without TROP2 sequence was used as control (Stuff). Human TROP2 cDNA was used to generate hTROP2 expressing plasmid (GeneCopoeia), and empty vector (EV) was used as control plasmid. Reconstituted cells were selected using hygromycin to get stably infected pools. Mouse TROP2 expression plasmids used in the rescue experiments all contain silence mutations at the sgRNA-targeting region.

### Immunoprecipitation and Immunoblotting

Immunoprecipitation was performed using Pierce Classic Magnetic IP Kit (Thermo Scientific, 88804) with minor modifications. Briefly, 1-3 million cells were lysed with IP lysis buffer including protease/phosphatase inhibitor cocktail (Cell Signaling Technology, 5872) for 10 mins. The lysates were centrifuged at 13,000g for 10 mins at 4 degrees to remove debris. The lysate was precleared with IgG for 3-4h at 4 degrees before the overnight incubation with human TROP2 antibody (Invitrogen, 14-6024-82) or mouse TROP2 (R&D, AF1122). The eluted proteins were analyzed by immunoblotting following standard SDS-PAGE. For regular immunoblotting, cultured cells were washed with ice-cold PBS, lysed with 1X SDS lysis buffer (GenScript, MB01015), and analyzed by standard SDS-PAGE. The primary antibodies include anti-mouse TROP2 (Invitrogen, MA5-29829), anti-human TROP2 (Abcam, ab214488), anti-CLAUDIN7 (Invitrogen, 34-9100), anti-VINCULIN (Cell Signaling Technology, 13901) and anti-ACTIN (Cell Signaling Technology, 3700).

### In Vivo Tumor Study and Treatment

All animal experiments were approved by the Institutional Animal Care and Use Committee at Mass General Hospital. Eight-week-old female BALB/C (Jackson Laboratory, 000651), NSG (Jackson Laboratory, 005557) and nude (MGH Cox-7 gnotobiotic animal facility) mice were used for in vivo studies. Mice were randomly allocated where appropriate. For antibody and drug treatments, tumors were first measured and randomized into each treatment group to control similar average starting size. 4T1 and AT-3 cells were injected in the mammary fat pad at 0.4- 1 × 10^5^ cells in 100 μl PBS, unless otherwise stated. Tumors were measured with calipers on the indicated days (volume = 0.5 × length × width^2^). Tumors were harvested for immunophenotyping by flow cytometry analysis, immunohistochemistry (IHC), or immunofluorescence (IF) at the experiment’s end point.

For anti-PD1 and hRS7 treatment experiments, 4T1 Trop2 KO+hTROP2 were injected into BALB/C mice and allowed to grow to reach approximately 100 mm^3^, the mice were then randomized and allocated to four treatment groups. Isotype control IgG (InVivomab, BE0089), anti-PD1 (InVivomab, BE0146) antibodies and/or hRS7 (Gilead) were injected intraperitoneally twice per week until the end point. The treatment doses were 250 μg/injection for SG and 100 μg/injection for IgG/anti-PD1. Tumor measurements were blinded whenever possible.

For CD8**^+^** T cell depletion, each mouse was administered intraperitoneally 200 µg anti-mouse CD8 (BioxCell, BE0061) or IgG2b isotype control (BioxCell, BE0090) one day before tumor inoculation and twice per week thereafter.

Doxycycline-inducible Claudin7 knockdown tumor cells (VectorBuilder, VB250922-1704szf) and control tumor cells (VectorBuilder, VB010000-9913att) were implanted into BALB/C mice. Seven days after tumor cell injection, tumors were measured and tumor-bearing mice were randomized and provided either doxycycline (2.0 mg/mL)-containing sucrose (2%) water or sucrose water alone continuously. Mice were subsequently treated with isotype control IgG or anti-PD-1 antibody twice weekly until reaching humane endpoints. Tumors were then harvested for flow cytometric analysis of tumor-infiltrating lymphocytes.

### Flow Cytometry Analysis

Single-cell suspensions of tumors were obtained by mincing and passing the tissues through 70 μm filters. Cells were stained with Viability Ghost Dye (Tonbo Biosciences, 13-0870) at 4 °C for 10 mins, followed by anti-CD16/32 blocking (Tonbo Bioscience, 70-0161-U500) at 4 °C for 10 min. For surface proteins, cells were stained with anti-CD45 (BioLegend, 103108), anti-CD3 (BioLegend, 100209), anti-CD8a (BioLegend, 100714), anti-TIM3 (BioLegend, 119703), anti-CD44 (BioLegend, 103041), anti-CD62L (BioLegend, 104410), and anti-PD1 (BD Biosciences, 563059) in FACS buffer (2% FBS in PBS). For cytokine staining, cells were treated with anti-CD3/CD28 (Thermo Fisher, 11452D) at 37 °C overnight and then with BD GolgiPlug (BD Biosciences, 550583) at 37 °C for 5 h. Following surface staining, cells were permeabilized using BD Cytofix/Cytoperm kit (BD Biosciences, 554714), and stained with anti-Granzyme B (Invitrogen, 48-8898-82). All antibodies were used at 1:150 dilution. Cells were fixed using 1% paraformaldehyde and analyzed by BD FACSAria. Data analysis was done on BD FACSDiva and FlowJo software.

For splenocytes, single cell suspension was obtained in the same manner as tumors. For 4T1 surface TROP2 staining, live cells were directly stained with anti-mouse TROP2 (R&D, FAB1122A).

### Immunofluorescence and Immunohistochemistry

Tumor tissues were carefully dissected and embedded in Tissue-Tek optimal cutting temperature compound (OCT, Sakura Finetek USA). 20um tissue section was collected and fixed in 4% PFA (diluted from 16% stock in PBS) for 15 minutes at room temperature. After washing with PBS for 3 times every 5 min to remove OCT and then incubate in blocking solution (5% normal donkey serum, 0.25% Triton X-100 in PBS) for 1h at room temperature, followed by incubation with anti-Claudin7 antibody (1:100 dilution; Invitrogen, 34-9100) and anti-Trop2 antibody (10ug/ul; RD system, AF1122) in blocking solution overnight at 4°C. After proper washing, Alexa Fluor 488-conjugated donkey anti-goat secondary antibody (1:500 diluted in blocking solution) for Trop2 and Alexa Fluor 568-conjugated donkey anti-rabbit secondary antibody (1:500 diluted in blocking solution) for Claudin 7 were applied to the tissue section for 1h at room temperature. After washing with PBS four times, tissue sections were preserved in mounting medium. The specimens were mounted with Vectashield and examined with a Zeiss Imager Z2 confocal microscope with 40× NA 1.40 oil differential interference contrast objective at room temperature (22°C) using Zeiss Zen 2.6 Blue edition software.

For immunohistochemistry, formalin-fixed paraffin-embedded (FFPE) mouse tumor tissues were cut into 4µm sections for staining. Samples were deparaffinized with xylene and then decloaked. Antibodies were incubated at room temperature for 30 min, followed by incubation with rabbit polymer for 30 mins. The samples were stained with 3,3′-Diaminobenzidine, counterstained with hematoxylin and then lithium carbonate, and dehydrated to xylene. Antibodies included anti-CD3 (Dako, A0452), anti-CD4 (Cell Signaling Technology, 25229), anti-CD8 (ebioscience, 14-0808-80), anti-4-1BB (Cell Signaling Technology, 18798) and anti-CD69 (ABclonal, A26620PM). Imaging of the samples was performed on OLYMPUS DP74 microscope. Percentages of CD3-positive cells were quantified using QuPath software (v.0.4.1) (https://qupath.github.io). Tumor/stromal interface for CD3^+^, CD4^+^ and CD8^+^ quantification was defined as a region spanning 250μm on either side of the tumor–stroma border. Staining of tumor Occludin used Occludin (E6B4R) Rabbit mAb (CST, 91131), and the analysis for signal intensity was done using Fiji/ImageJ software (https://fiji.sc/).

For Claudin7 immunofluorescence in Trop2 KO 4T1 rescued with full length or ECDTM TROP2, the staining protocol of FFPE samples was identical. The samples were incubated with anti-Claudin7 antibody (1:50 dilution, Invitrogen, 34-9100) at room temperature for 1h, followed by Alexa Flour 555-conjugated goat ant-rabbit secondary antibody (1:500 dilution in blocking buffer). Imaging of the samples was done on Zeiss Imager Z2 confocal microscope. Quantitation of immunofluorescence was performed using HALO image analysis platform (Indica Labs). Using the Area Quantification FL (v2.3.4) module, intensity thresholds were set globally across all 20x confocal images to identify pixels positive for Claudin7 Alex Fluor 568 fluorescent stain. The percentage of positive pixels out of all pixels within the manually annotated tumor area was determined for each image. The images were manually annotated to exclude staining and imaging artifacts from analysis. Cell segmentation and compartment quantification were performed using the HALO HighPlex FL module, which included nuclear detection with the default AI algorithm based on DAPI signal. Cell compartments were assigned as “membrane” and “non-membrane”. For each cell, the average intensity of Claudin7 Alexa Fluor 568 signal per compartment was determined. Due to the expected membranous staining of Claudin7, a positivity threshold for each image was set as the median non-membrane signal.

Immunofluorescence apoptosis TUNEL staining was completed by iHisto Inc. Briefly FFPE slides were baked overnight at 60°C and deparaffinized and rehydrated. Slides were washed in PBS, PBS, and PBST for 5 minutes each, respectively. Proteinase K working solution was applied to each section and sections were allowed to incubate at 37°C for 20 minutes. TdT Equilibration Buffer was applied to each sample and incubated at 37°C for 10 minutes. Excess TdT Equilibration Buffer was removed from the sections and the Labeling Working Solution was applied to each section. Slides were incubated at 37°C for 2 hours and washed as before. Sections were stained with DAPI and coverslipped using Fluoroshield.

### RNA-Seq

RNA sequencing was done using 4T1 *Trop2* WT and KO tumors frozen sections. Total RNA was extracted using RNeasy Plus Mini Kit (Qiagen, 74134). RNA integrity was checked with Agilent 4200 TapeStation System. Following DNA contaminant removal and rRNA depletion, RNA-Seq libraries were constructed using NEBNext Ultra II RNA Library Preparation Kit for Illumina. In brief, RNAs were fragmented at 94°C for 15 min. First and second strand cDNA synthesis was done using primers for random priming sites and Unique Molecular Identifiers (UMIs), which were incorporated into final cDNA. After adapter ligation, cDNAs were purified and enriched by PCR to generate sequencing libraries. RNA-Seq libraries were sequenced using Illumina HiSeq 400 platform, and analysis was done on HiSeq Control Software (HCS). Raw fastq file for each tumor sample was generated by demultiplexing with Illumina bcl2fastq 2.17 software. Gene set enrichment analysis was done using GSEA 4.3.2 software (gsea-msigdb.org).

### Bioinformatic analysis

For single cell data processing, original data were retrieved from European Genome-phenome Archive (EGA) and Gene Expression Omnibus (GEO) with data accession no. EGAD00001006608 (30). SCTransform was used to normalize the data. AggregateData function in muscat package was used to generate pseudobulk counts for cancer epithelial cells and total immune cells. EdgeR package was used to perform TMM standardization after 0 was removed to generate counts per million (CPM) expression matrix. For bulk RNA-seq analysis in GSE241876 (31), same edgeR based methods as mentioned above for single cell were used to generate CPM expression matrix. For the calculation of Odds Ratios, samples were divided into two groups according to the CPM values of the gene. The OptimalCutpoints package was used to calculate the Youden index to determine the optimal cutoff value, and the logistic regression was then used to generate OR and 95% CI. Non-responsive group (NR) includes progressive disease (PD) and stable disease (SD), and responsive group (R) includes partial responses (PR) and complete responses (CR). T cell receptor clonotype expansion or non-expansion categories were defined by original paper through single-cell T cell receptor (TCR)-seq.

## Acknowledgements

We gratefully acknowledge Immunomedics/Gilead for the gift of hRS7. We thank Grace Dejun Wu, Akiko Suzuki, David Li and MGH Histopathology Research Core for their assistance. Ting Liu acknowledges salary support from the National Natural Science Foundation of China (No. 82303702).

## Ethics approval and consent to participate

All animal experiments were approved by the Institutional Animal Care and Use Committee at Mass General Hospital (protocol: 2004N000228). All patients provided written informed consent under Institutional Review Board-approved protocols at Mass General Hospital (protocol: 2018P001865).

## Consent for publication

All authors approve the manuscript for publication.

## Availability of data and material

Experimental materials, raw data supporting the figures and graphs as well as full blots are available from the authors upon request.

## Authors’ contributions

B.W. and L.W.E. conceived, designed, and supervised the project and wrote the initial manuscript. B.W., W.T., E.B., J.L., B.K.P., K.H.X., B.G., Y.C., K.J., and F.S. performed the experiments. T.L., E.I.P., C.N., L.T.N., D.T.T., N.T., S.S., R.O.A., V.I.B., S.J.I., L.M.S., and A.B. contributed to data analysis, manuscript review, and editing. L.W.E. is responsible for the overall content as guarantor. All authors approved the final manuscript.

