## Supplemental for "A TROP2/Claudin Program Mediates Immune Exclusion to Impede Checkpoint Blockade in Breast Cancer"

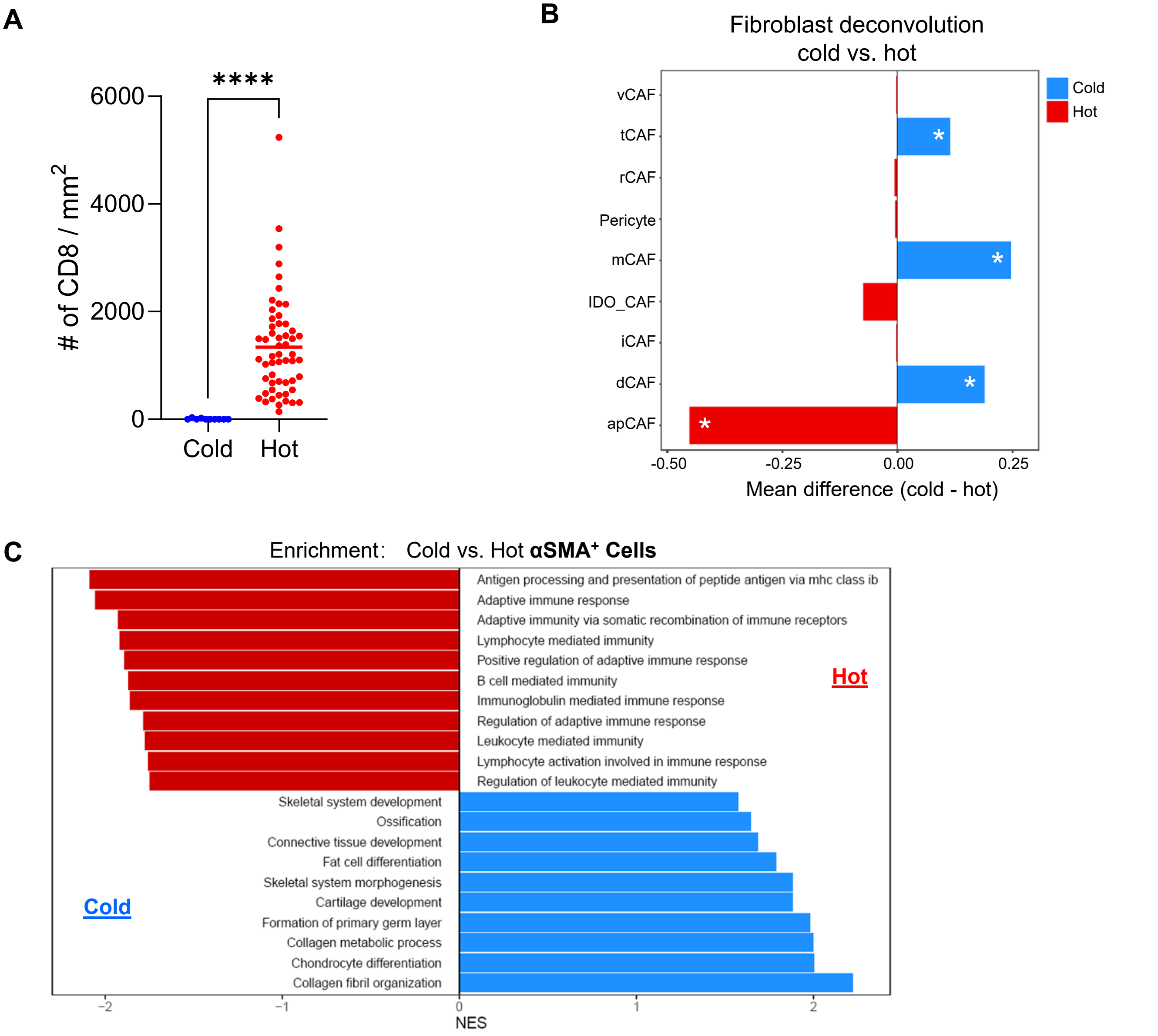


**Supplementary Figure S1. Spatial transcriptomics identifies fibroblast specific programs associated with TNBC immune phenotypes. A,** $CD8^+$ cell density (cells/mm²) in immune-cold versus immune-hot ROIs. **B,** Computational deconvolution of $\alpha SMA^+$ cancer associated fibroblast (CAF) subsets in immune-cold versus immune-hot ROIs. vCAF: vascular CAF, tCAF: tumor-like CAF, rCAF: reticular-like CAF, mCAF: matrix CAF, iCAF: inflammatory CAF, dCAF: dividing CAF, apCAF: antigen-presenting CAF, *$padj$<0.05. **C,** Top 10 pathways associated with $\alpha SMA^+$ cells in immune-cold versus immune-hot ROIs.

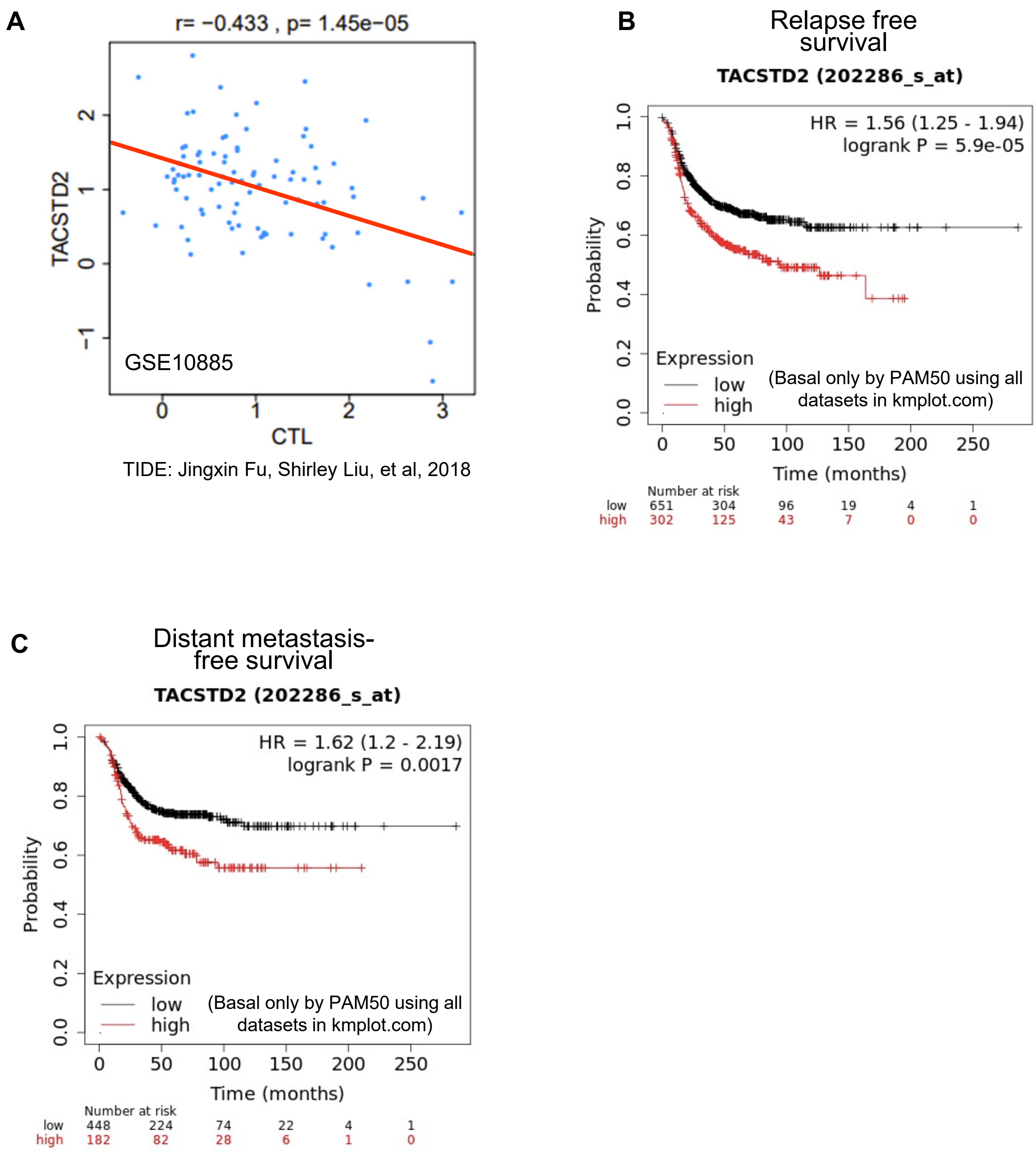


**Supplementary Figure S2. High TROP2 associates with cold TIME and worse survivals. A,** Correlation between TROP2 and cytotoxic T lymphocyte score (CTL) in GSE10885 dataset by TIDE analysis. **B** and **C,** Correlation between TROP2 and relapse free survival (B) or distant metastasis free survival (C) in basal subtype of breast cancer, generated by Kaplan-Meier Plotter database (https://kmplot.com/analysis/).

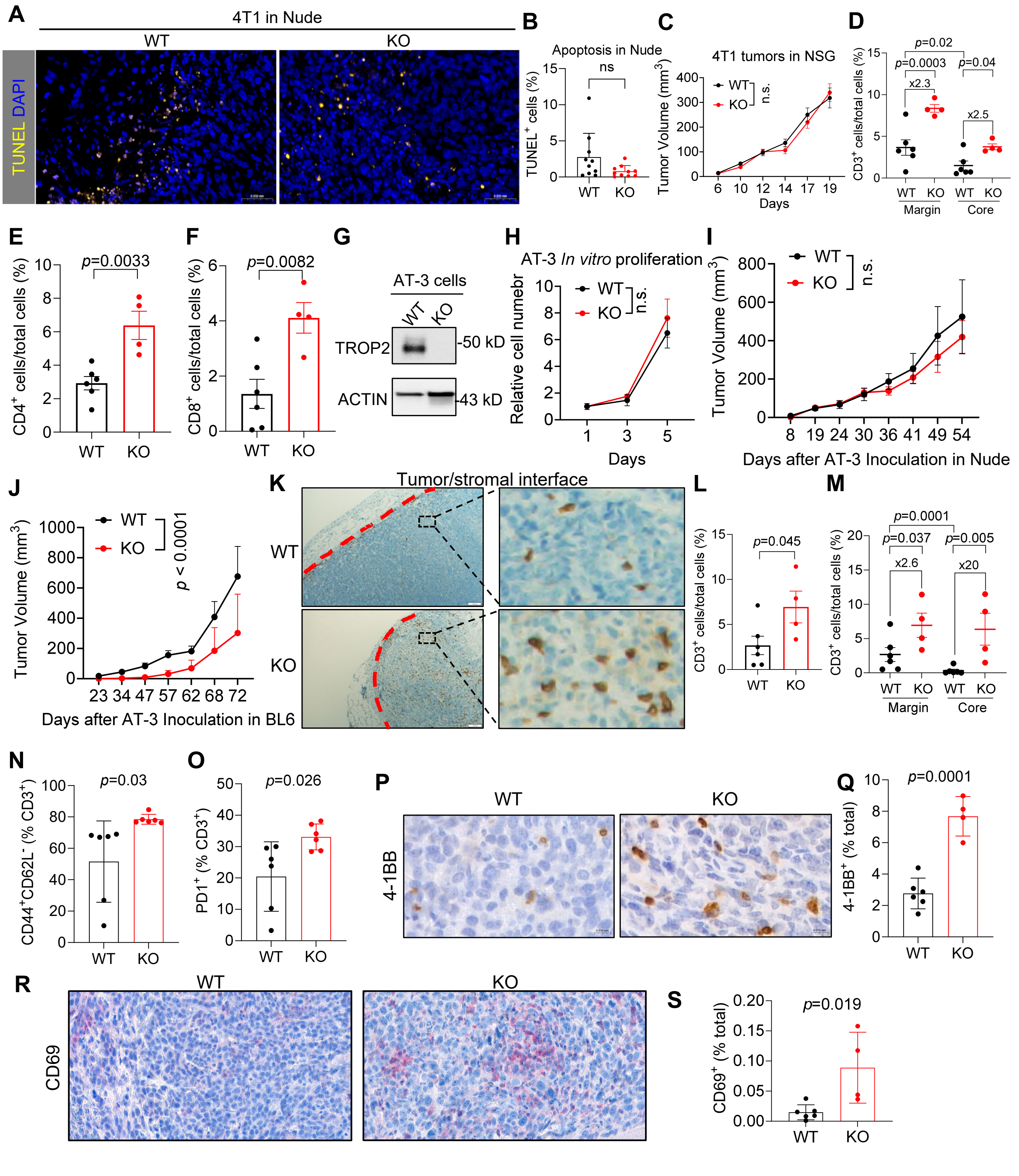


**Supplementary Figure S3. TROP2 promotes mammary tumor growth and immune exclusion in immunocompetent hosts. A** and **B,** Representative images (A) and quantification (B) for TUNEL apoptosis staining for wildtype (WT) and *Trop-2* knockout (KO) 4T1 tumors harvested from nude mice. **C,** WT and KO 4T1 tumor growth in immunocompromised (NSG) hosts (n = 10 tumors per group). **D,** Comparison of $CD3^+$ T cells in the margin and core of 4T1 tumors from the immunocompetent hosts. **E** and **F**, Quantification of $CD4^+$ and $CD8^+$ T cell at tumor/stromal interface by IHC staining in WT and KO 4T1 tumors. **G,** Western blot for wildtype (WT) and *Trop-2* knockout (KO) AT-3 cells. **H-J,** *Trop2*-WT and -KO AT-3 tumor growth in cell culture (H), in immunocompromised (nude) hosts (WT, n = 4 tumors; KO, n = 3 tumors) (I), and in immunocompetent C57BL/6 (BL6) hosts (WT, n = 6 tumors; KO, n = 4 tumors) (J). **K** and **L,** Representative images (K) and quantification (L) of $CD3^+$ T cell IHC staining of WT and KO AT-3 tumors. **M,** Comparison of $CD3^+$ T cells in the margin and core of AT-3 tumors. **N** and **O,** $CD44^+CD62L^-$ effector memory (N) and $PD1^+$ activated (O) $CD3^+$ T cells in WT and KO 4T1 tumors. **P** and **Q,** Representative images (P) and quantification (Q) of T cell activation marker 4-1BB IHC staining of WT and KO 4T1 tumors. **R** and **S,** Representative images (R) and quantification (S) of T cell activation marker CD69 IHC staining of WT and KO 4T1 tumors.

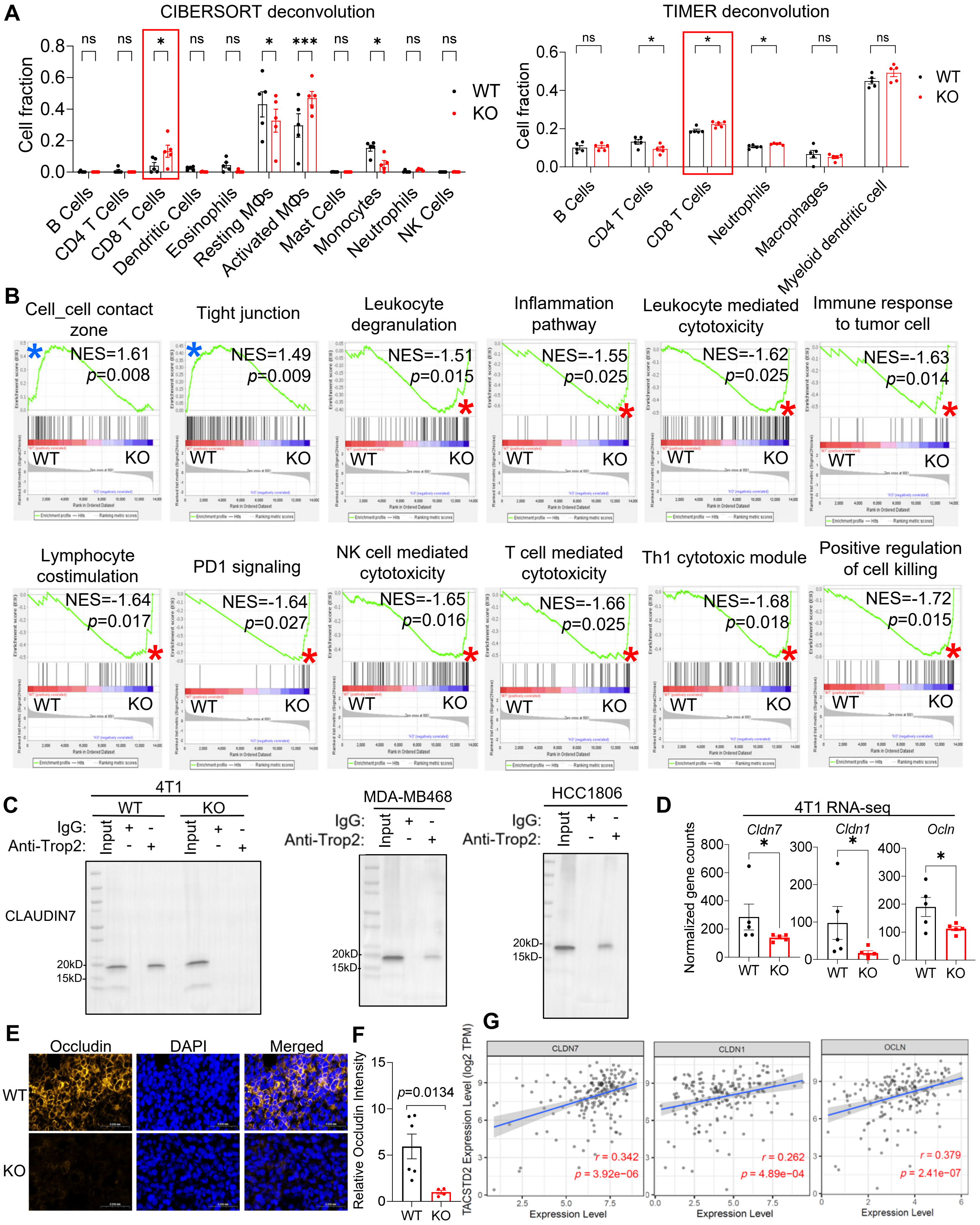


**Supplementary Figure S4. RNA-seq analysis and Occludin staining for WT and KO 4T1 tumors. A,** Computational deconvolution of tumor-infiltrating immune subsets from bulk RNAseq of WT and KO 4T1 tumors using CIBERSORT (left panel) and TIMER algorithm (right panel). Red boxes highlight significant differences in CD8+ T cell populations. **B,** Gene set enrichment analysis of bulk RNA-seq from *Trop2*-WT and -KO 4T1 tumors (blue asterisk: positive NES; red asterisk: negative NES). **C,** Full membrane of CLAUDIN7 IP-WB from Fig. 4B. **D,** Gene expression (normalized gene counts) for Claudin-7 (*Cldn7*), Claudin-1 (*Cldn1*), and Occludin (*Ocln*) from RNAseq of WT and KO tumors. **E,** Representative tumor tissue immunofluorescence images of Occludin and DAPI (nuclear stain) showing loss of Occludin expression with TROP2 knockout. **F,** Quantification of Occludin intensity in WT and KO 4T1 tumors for panel E. **G,** Correlation between TROP2 and CLDN1/CLDN7/OCLN expression in basal breast cancer patient cohorts by TIMER analysis using TCGA database.

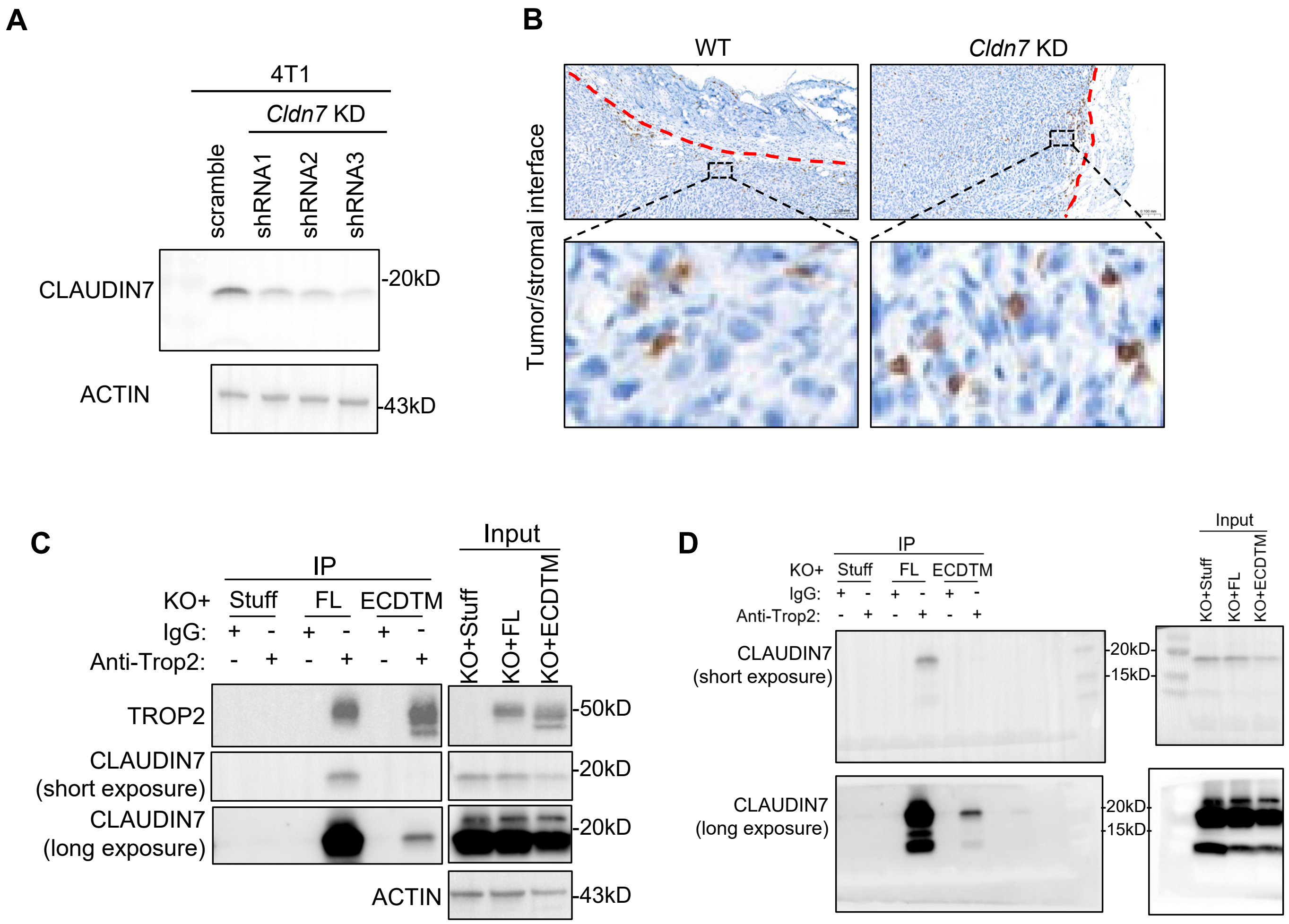


**Supplementary Figure S5. Cldn7 knockdown and interaction between truncated TROP2 and CLAUDIN7. A,** Western blot for wildtype (WT) and *Cldn7*-knock down (KD) 4T1 cells. **B,** Representative images of $CD3^{+}$ T cell IHC staining of *Cldn7*-WT and -KD 4T1 tumors from BALB/c mice. **C,** Western blot showing co-IP of TROP2 and Claudin 7 using IgG and anti-TROP2 in *Trop2*-KO 4T1 reconstituted with Stuff, Full-Length TROP2, or ECDTM truncated mutant. **D,** Full membrane of panel C. FL: full length.

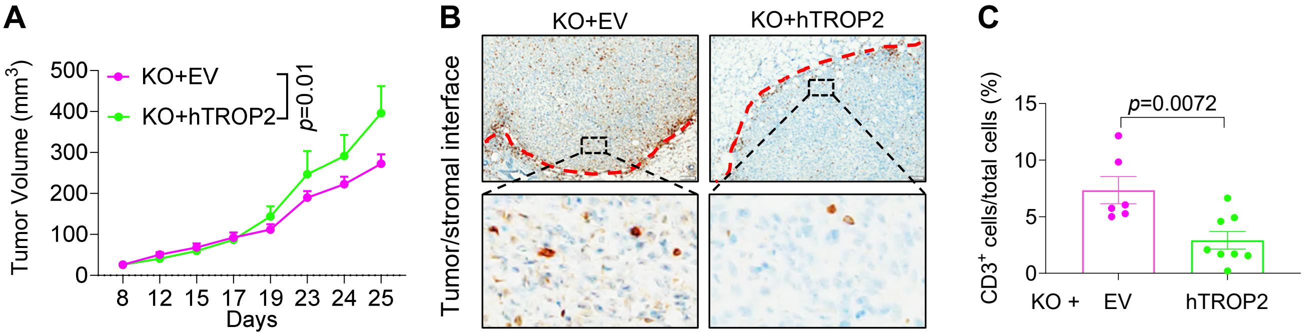


**Supplementary Figure S6. Human TROP2 reconstitution in 4T1 *Trop2*-KO tumor. A,** Tumor growth curves for KO+EV and KO+hTROP2 4T1 in BALB/C mice (KO+EV, n = 6 tumors; KO+hTROP2, n = 8 tumors). **B** and **C,** Representative images (B) and quantification (C) of $CD3^{+}$ T cell IHC staining of KO+EV and KO+hTROP2 tumors.

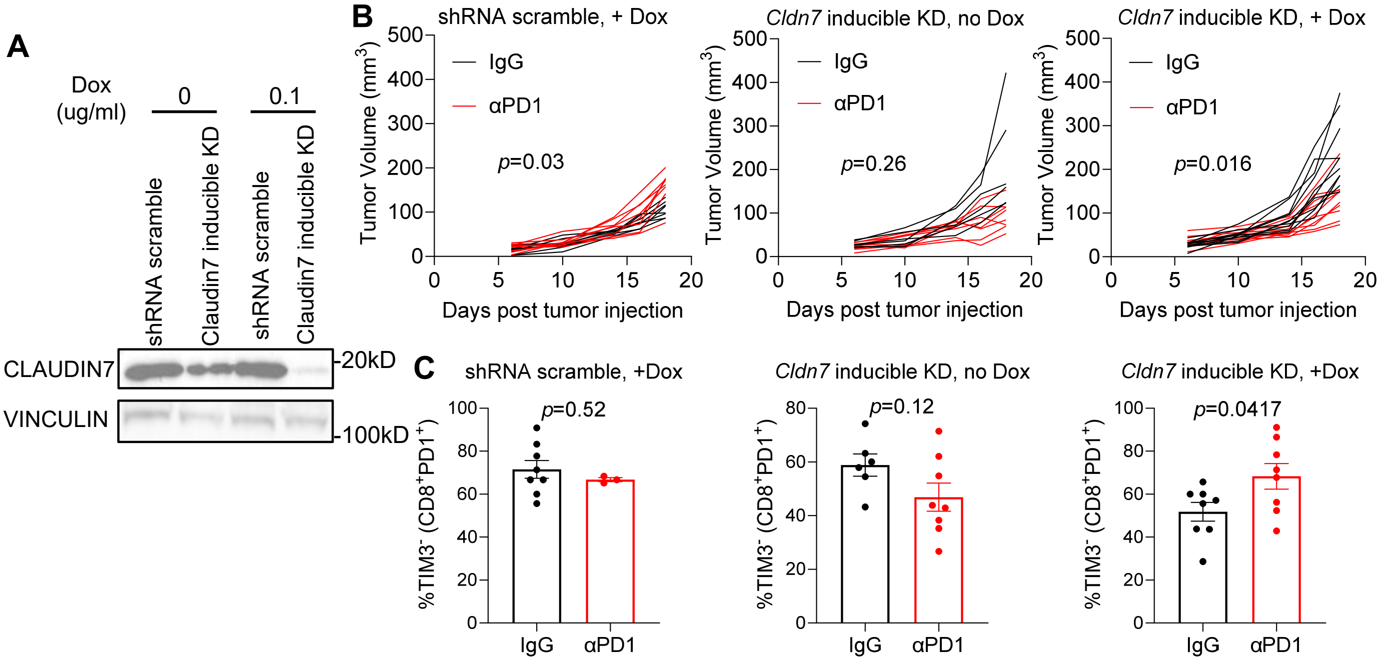


**Supplementary Figure S7. Caludin7 knockdown sensitizes 4T1 tumors to anti-PD1. A,** Western blot of doxycycline (Dox) inducible *Claudin7* knockdown (KD) in 4T1 cells. **B,** Tumor growth curves for shRNA scramble or *Cldn7* inducible KD 4T1 in BALB/C mice with or without Dox and with or without IgG or anti-PD1 treatment. **C,** Quantification of $TIM3^-$ early activated $CD8^+PD1^+$ T cells in tumors harvested from panel B.

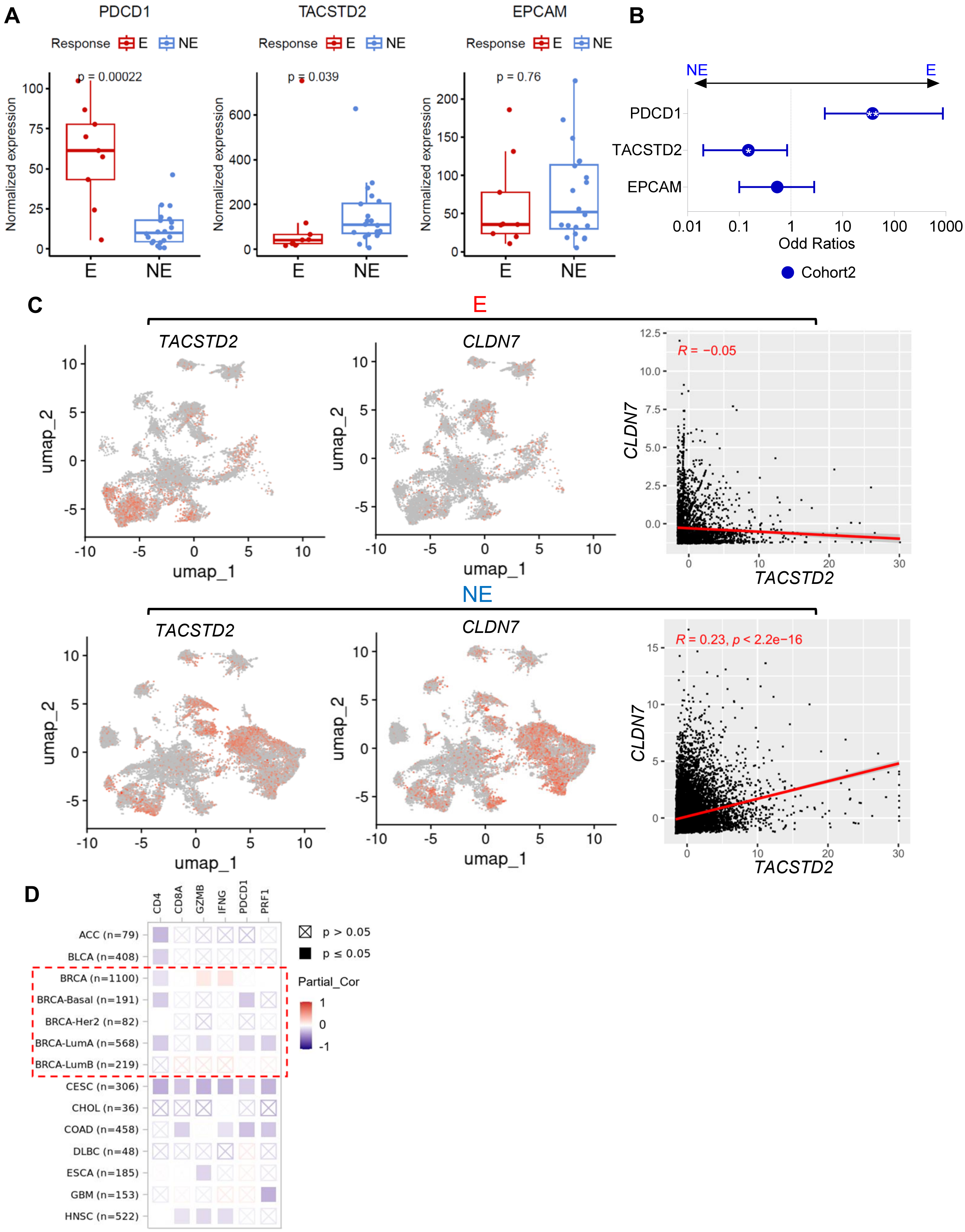


**Supplementary Figure S8. Correlations between TACSTD2 and immune checkpoint inhibitor response. A,** PDCD1 (PD1), TACSTD2 (TROP2) and EPCAM expression derived from pseudobulk analysis of single-cell RNAseq data in TCR clonotype expanded (E) vs. non-expanded (NE) cases following pre-operative single-agent pembrolizumab (30). **B,** Odds ratios for associations between gene expression within the epithelial cell populations and responses (E vs. NE) in the Cohort in (A). **p*<0.05, ***p*<0.01. **C,** UMAPs and correlation of *TACSTD2* and *CLDN7* expression in the tumor epithelial cells from E (upper panel) and NE (lower panel) cases in the Cohort in (A). **D,** Correlation between *EPCAM* and immune cell markers in multiple cancer types by TIMER analysis in the TCGA database. Red dashed box indicates breast cancers.
